# Antagonistic functions of the two oxidative pentose phosphate pathway dehydrogenases in shaping oxidative stress responses

**DOI:** 10.64898/2026.05.01.722205

**Authors:** Lug Trémulot, Zheng Yang, Gilles Châtel-Innocenti, Patrick Willems, Katrien Van Der Kelen, Hélène Vanacker, Emmanuelle Issakidis-Bourguet, Frank Van Breusegem, Amna Mhamdi, Graham Noctor

## Abstract

The oxidative pentose phosphate pathway (OPPP) is a source of cellular NADPH, generated through the sequential activities of glucose-6-phosphate dehydrogenase (G6PDH) and 6-phosphogluconate dehydrogenase (6PGDH). Using the catalase-deficient *cat2* background as a model for H_2_O_2_-triggered salicylic acid (SA) signaling we identified the cytosolic G6PDH isoform G6PD5 as a key determinant of redox homeostasis and SA-dependent defense activation (Trémulot et al., companion manuscript). However, the mechanisms underlying this function remain enigmatic. In this work, genetic and transcriptomic analyses show that the role of G6PD5 cannot be explained solely by altered NADPH generation for either NADPH oxidases or the ascorbate-glutathione pathway, suggesting other possible links. To identify such links, a forward genetic screen was employed. We searched for mutations that modulate the suppressed lesion phenotype in *cat2 g6pd5* in a photorespiration-dependent manner. This screen identified a mutation in *PGD2*, encoding the cytosolic 6PGDH. Strikingly, functional analyses of mutants and overexpression lines revealed that PGD2 exerts effects opposite to those of G6PD5 in SA signaling. Our observations uncover an unexpected antagonism between the two cytosolic NADPH-producing steps within the OPPP. Pharmacological analyses support a signaling role for the metabolic intermediate 6-phosphogluconolactone in linking the OPPP to SA signaling. These findings indicate that the OPPP is not solely a source of reducing power during oxidative stress but also acts as a signaling module in which metabolic intermediates contribute to the control of stress-induced immune responses.

## Introduction

Oxidative stress is an important feature of many plant responses to the environment, and occurs when cells shift to a more oxidized state, notably due to the production of reactive oxygen species (ROS) (Halliwell, 2006; Mittler, 2017). This effect triggers signaling cascades, enabling plant cells to respond appropriately to stresses such as pathogen attack (Herrera-Vásquez et al., 2015; Mittler, 2017). The connection between oxidative stress and defense signaling is well-established. For example, in the Arabidopsis *cat2* mutant, in which catalase activity is decreased by about 80%, oxidative stress activates salicylic acid (SA)-dependent accumulation and signaling and promotes the formation of hypersensitive response (HR)-like lesions (Chaouch et al., 2010). A key advantage of the *cat2* system is its conditionality. Since the main role of CAT2 is to remove photorespiratory hydrogen peroxide (H_2_O_2_), the intensity of oxidative stress in *cat2* can be easily manipulated by altering external conditions such as irradiance and CO_2_ levels (Queval et al., 2007). Hence, while oxidative stress, leaf lesions and associated defense signaling are observed in *cat2* grown in standard conditions (ambient air), all these phenomena are suppressed when the mutant is grown at high CO_2_ (hCO_2_) levels that largely annul photorespiration.

NADPH is a pyridine dinucleotide that participates in metabolism and redox homeostasis in numerous ways. Within the context of oxidative stress, NADPH sustains ROS production by Respiratory Burst Oxidase Homologs (RBOHs), supports peroxidase-based H_2_O_2_ removal, notably via the ascorbate-glutathione pathway, and enables thiol-based signaling (Rouhier, 2010; Dietz, 2011; Foyer and Noctor, 2011; Geigenberger et al., 2017; Kaya et al., 2019; Noctor, 2025). All of these processes could be important in the *cat2* system undergoing oxidative stress. Indeed, a previous study has shown that *cat2*-triggered metabolic responses can be to some extent modulated by *AtRBOHF*, encoding one of the two major leaf NADPH oxidases (Chaouch et al., 2012). However, NADPH generation for antioxidant functions is expected to be particularly important in *cat2*. Since catalase itself does not require reductants, loss of its function in standard growth conditions (air) imposes a greater load on NADPH-dependent pathways to metabolize H_2_O_2_, a phenomenon reflected by up-regulation of enzymes of the ascorbate-glutathione pathway and oxidation of glutathione (Tuzet et al., 2019; Noctor, 2025).

NADPH regeneration therefore becomes particularly important during oxidative stress and could play a role in the transmission of redox signals. However, the relative importance of specific dehydrogenases for the various NADPH-dependent processes associated with oxidative stress remains unclear. Previous investigations have explored the importance of specific cytosolic NADPH-generating enzymes using Arabidopsis mutants. For instance, several studies have focused on the roles of isocitrate dehydrogenase and malic enzyme (cICDH and NADP-ME) in responses to pathogens and oxidative stress (Mhamdi et al., 2010b; Voll et al., 2012; Li et al., 2013; Singh et al., 2016; Choudhury et al., 2018; Dangol et al., 2019; Wu et al., 2022). The contribution of the two dehydrogenases of the oxidative pentose phosphate pathway (OPPP) in the signaling triggered by oxidative stress is less studied.

The OPPP is the initial part of the pentose phosphate pathway and involves three steps, two of which generate NADPH. Through these three steps, glucose 6-phosphate (G6P) is converted to ribulose-5-phosphate and CO_2_, generating NADPH from NADP_+_ in steps catalyzed by glucose-6-phosphate dehydrogenase (G6PDH) and 6-phosphogluconate dehydrogenase (6PGDH) (Kruger and von Schaewen, 2003). In plants, genetic evidence linking the OPPP to oxidative stress responses is limited, although some previous work has explored the influence of G6PDH in biotic stress (Scharte et al., 2009; Stampfl et al., 2016; Hu et al., 2019). One notion is that cytosolic G6PDH could be particularly important in generating NADPH for RBOH-dependent ROS signaling (Scharte et al., 2009). In terms of antioxidant-related functions, modeling suggests that during oxidative stress the OPPP can account for a large part of the regeneration of the required NADPH (Tuzet et al., 2019).

In the preceding study (Trémulot et al., companion article), we identified cytosolic G6PD5 as a key determinant of redox homeostasis during oxidative stress. Loss of *G6PD5* function also fully abolished oxidative stress-triggered SA signaling, establishing a functional link between cytosolic OPPP activity and activation of hormonal and defense reactions. These findings highlight a role for G6PD5 in the transmission of redox-derived signals. In the present study, we address the mechanisms through which this occurs. Specifically, we used comparative transcriptomics to indicate whether the effect of loss of *G6PD5* function might largely be explainable by compromised activities of key NADPH-dependent pathways and, if not, whether other mechanisms could be in play. Because this first analysis provided little evidence that the effect of the *g6pd5* mutation can be explained by compromised NADPH-requiring pathways, we conducted a two-step forward genetic screen for mutants that revert *cat2 g6pd5* to the *cat2* phenotype in a photorespiration-dependent manner. This approach identified *PGD2*, encoding the cytosolic 6PGDH, as a modulator of oxidative stress phenotypes and signaling. Further analyses revealed that *G6PD5* and *PGD2* act antagonistically in influencing oxidative stress responses and suggest that this might reflect a signaling role for a metabolic intermediate of the cytosolic OPPP.

## Results

### Exploring *G6PD5* function through NADPH-dependent pathways

We reasoned that if alteration of NADPH supply for either RBOHs or the ascorbate-glutathione pathway makes an important contribution to the effect of the *g6pd5* mutation, then mutants impaired in these processes in the *cat2* background should display transcriptomic profiles similar to those observed in *cat2 g6pd5* (Fig. 1A). As previously reported, *cat2 g6pd5* exhibits strong transcriptomic similarity with *cat2 sid2*, a genotype defective in SA biosynthesis (Supplementary Figure S1; Trémulot et al., companion article), thereby providing a reference signature for suppression of oxidative stress-triggered SA signaling. As a first step, we generated transcriptomic datasets for *cat2 atrbohd* and *cat2 atrbohf*, mutated in CAT2 and the two major RBOH isoforms expressed in Arabidopsis leaves (Chaouch et al., 2012), grown in the same conditions as *cat2 g6pd5* (Fig. 1; Experiment 1). In a second analysis, transcriptomes of *cat2 mdar2* and *cat2 dhar123*, impaired in enzymes of the ascorbate-glutathione pathway (Rahantaniaina et al., 2017; Xu et al., 2025), were compared for overlap with *cat2 g6pd5* (Fig 1; Experiment 2). Apart from the *cat2 mdar2* transcriptome, which has already been partly reported (Xu et al., 2025), these datasets were generated specifically for the present study.

**Figure 1.**
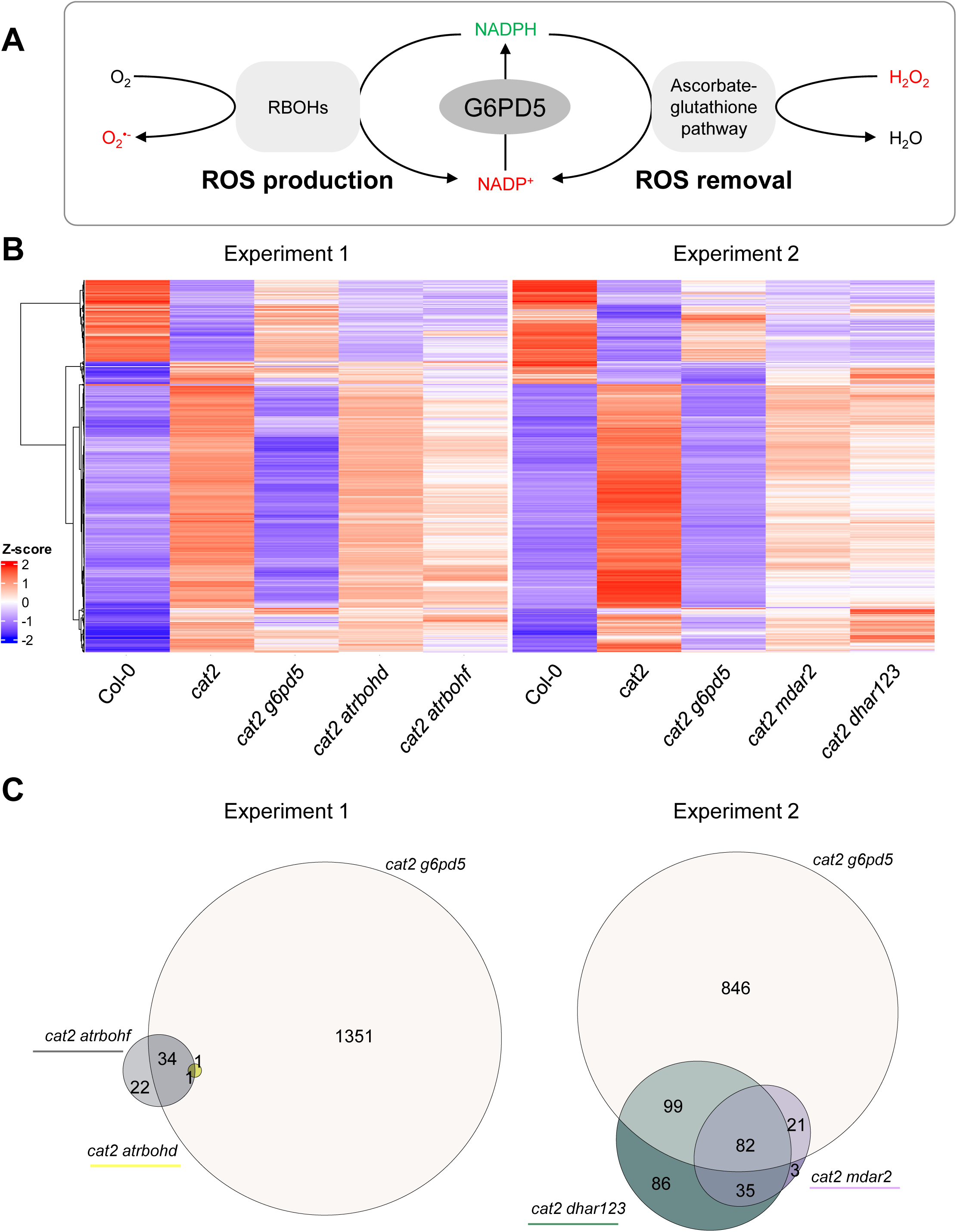
Comparison of transcriptomic signatures in *cat2 g6pd5* and mutants impaired in NADPH-dependent pathways. A. Scheme illustrating possible roles of G6PD5-derived NADPH in oxidative stress signaling. NADPH may supply RBOH, NADPH oxidases, promoting ROS production, or ascorbate-glutathione pathways, which contribute to ROS processing. B. Heatmap of gene expression profiles in Col-0, *cat2*, *cat2 g6pd5* and the mutants defective in NADPH utilizing pathways. Two independent transcriptomic datasets were analyzed: experiment 1 (*cat2 atrbohf* and *cat2 atrbohd*) and experiment 2 (*cat2 mdar2* and *cat2 dhar123*). The heatmap displays the 734 genes differentially expressed between Col-0 and *cat2* (DEGs) that are common to both experiments. Expression values are shown as Z-scores, calculated independently for each dataset. Hierarchical clustering was performed using Pearson correlation and Ward.D2 distance. C. Venn diagrams showing overlap of DEGs for the three contrasts comparing *cat2* with *cat2 g6pd5* or mutants related to NADPH consumption. The figures indicate the number of DEGs within each specific set. For instance, compared to the contrast considered for the experiment 2 dataset, 846 DEGs are specific for the *cat2 g6pd5* vs *cat2* comparison.

The transcriptomic profiles of the 734 genes that were differentially expressed (DEGs) in *cat2* relative to Col-0 in both experiments were examined in *cat2 atrbohd*, *cat2 atrbohf*, *cat2 mdar2*, and *cat2 dhar123*. In all four genotypes, expression patterns were more similar to *cat2* than to *cat2 g6pd5* (Fig. 1B). Mutations affecting the ascorbate-glutathione pathway slightly attenuated the *cat2* transcriptomic signature, but mutations affecting RBOHs had only a very weak effect. In all cases, the similarity with *cat2 g6pd5* remained limited. Overlap analysis showed that only 10% and 17% of the genes altered in *cat2 g6pd5* were shared with *cat2 mdar2* and *cat2 dhar123*, respectively (Fig. 1C, right panel). In contrast, strong similarity was observed in both experiments between *cat2 g6pd5* and *cat2 sid2*, a mutant impaired in SA biosynthesis (see also Trémulot et al., companion manuscript). In both transcriptomic datasets, the *cat2 sid2* and *cat2 g6pd5* profiles more closely resembled those of Col-0 rather than *cat2*, and approximately 60% of the genes differentially expressed in *cat2 g6pd5* vs *cat2* were also detected in the *cat2 sid2* vs *cat2* comparison (Supplementary Figure S1).

These analyses show that impairment of RBOHs or components of the ascorbate-glutathione pathway in the *cat2* background does not reproduce the transcriptional changes observed in *cat2 g6pd5*. Although G6PD5 contributes to cellular redox homeostasis through NADPH regeneration (Trémulot et al. companion manuscript), the transcriptomic analyses indicate that altered NADPH supply to these pathways is not sufficient to explain the abolished SA signaling in *cat2 g6pd5*. We therefore undertook a classical genetic screen to attempt to identify genes and pathways that could explain this effect.

### A genetic screen identifies *PGD2* as a candidate modulating SA signaling downstream of G6PD5

We conducted a double forward genetic screen in *cat2 g6pd5* (Fig. 2A). A first screen sought to identify M2 plants presenting a *cat2*-like lesion phenotype. In a second screen, we examined whether or not growth in hCO_2_ suppressed the lesion phenotypes in the F3 progeny of the backcrossed revertant lines (Fig. 2B). This strategy was chosen because growth at hCO_2_ abolishes photorespiratory H_2_O_2_ production and attenuates oxidative stress signaling and related phenotypes in *cat2* (Fig. 2; Queval et al., 2007). Hence, the second screen was designed to discriminate between lesions that are dependent on loss of *CAT2* function and those resulting from independent lesion-mimic mutations (Lorrain et al., 2003; Bruggeman et al., 2015; Janda and Ruelland, 2015). According to this logic, *cat2*-dependent lesions should be present in air but not hCO_2_, and hCO_2_ should attenuate phenotypes less strongly in mutants that show lesions independent of *cat2*-triggered oxidative stress.

**Figure 2.**
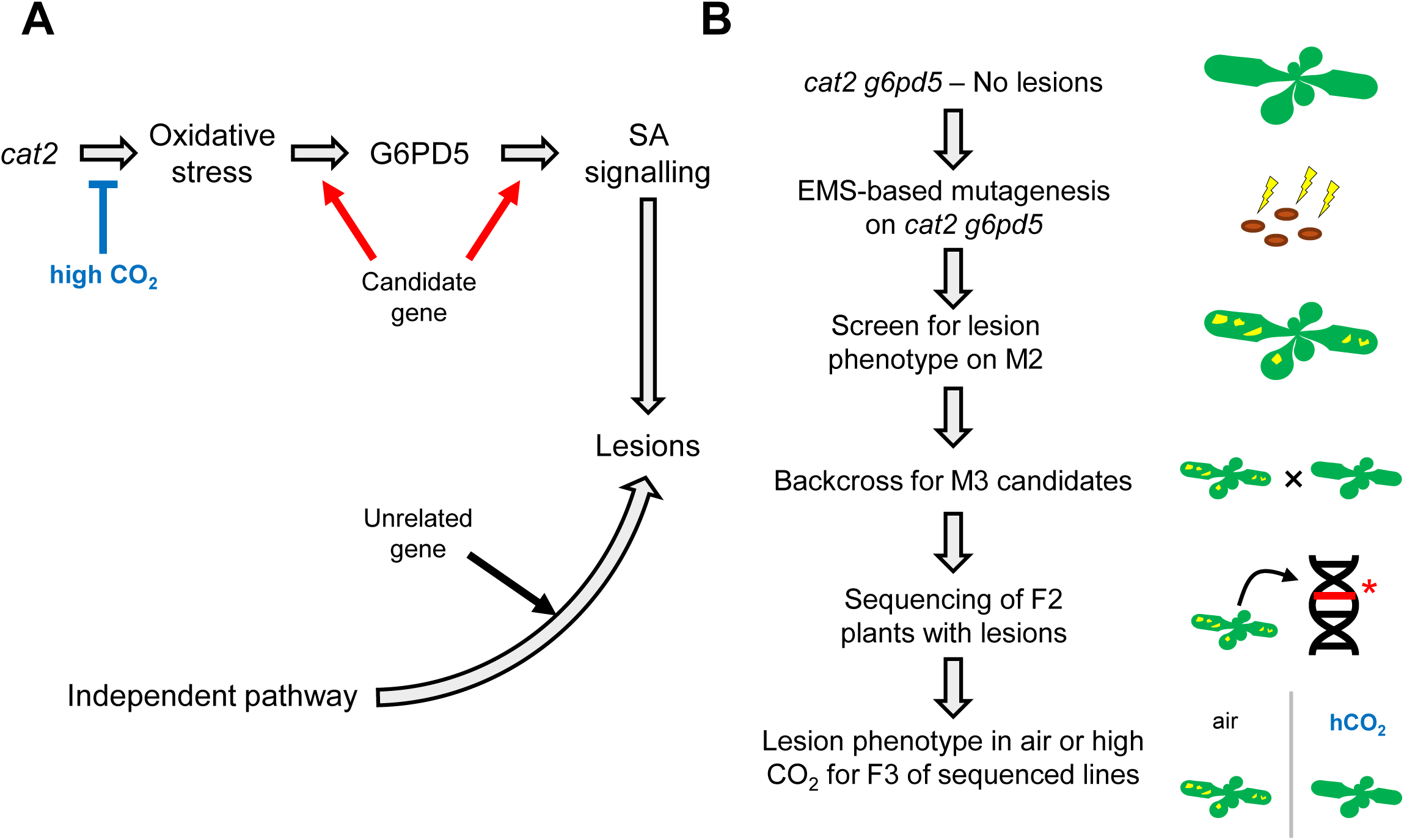
Strategy for a double genetic screen to identify suppressors of the *cat2 g6pd5* phenotype. A. Schematic representation of the screening strategy. Seeds of the *cat2 g6pd5* line were mutagenized and screened in the M2 generation for the reappearance of lesion formation. High CO_2_ conditions (hCO_2_) were used as a secondary filter to distinguish lesions dependent on the *cat2* oxidative stress background, as growth at elevated CO_2_ suppresses photorespiratory H_2_O_2_ production and associated lesion formation. B. Illustration of the screening workflow showing the successive steps of mutagenesis, identification of revertant lines displaying lesion formation, and verification of the hCO_2_ dependent phenotype through growth under ambient air or hCO_2_ conditions. The selected revertant lines (showing lesions) are backcrossed with *cat2 g6pd5,* and the F2 showing lesions are sequenced to identify the causal gene. Lines in which lesions were suppressed under CO_2_ were retained for subsequent analysis.

The primary screen of approximately 70 000 M2 plants grown in air identified several that presented *cat2*-like lesions in the *cat2 g6pd5* background. Segregation analysis of F1 and F2 progeny indicated that for five of these, the phenotype resulted from a recessive mutation at a single locus. These lines were subjected to a secondary screen to establish whether the lesions could be abolished at hCO_2_, which annuls *cat2*-triggered oxidative stress. Only one of the five lines (*cat2 g6pd5 17.21*) met this criterion (Table 1, Fig. 3A, Supplementary Figure S2). In the other four lines, lesion formation was either little affected (or else aggravated) by growth at hCO_2_, suggesting that in these mutants lesion development was not favored by photorespiration and was therefore not a consequence of *cat2*-triggered oxidative stress (Table 1, Supplementary Figure S2). Whole-genome sequencing of the four lines revealed non-synonymous mutations in genes previously associated with lesion phenotypes (Bowling et al., 1997; Meng et al., 2009). These were *MIPS1* (three allelic mutants) and *CPR5* (Table 1).

**Figure 3.**
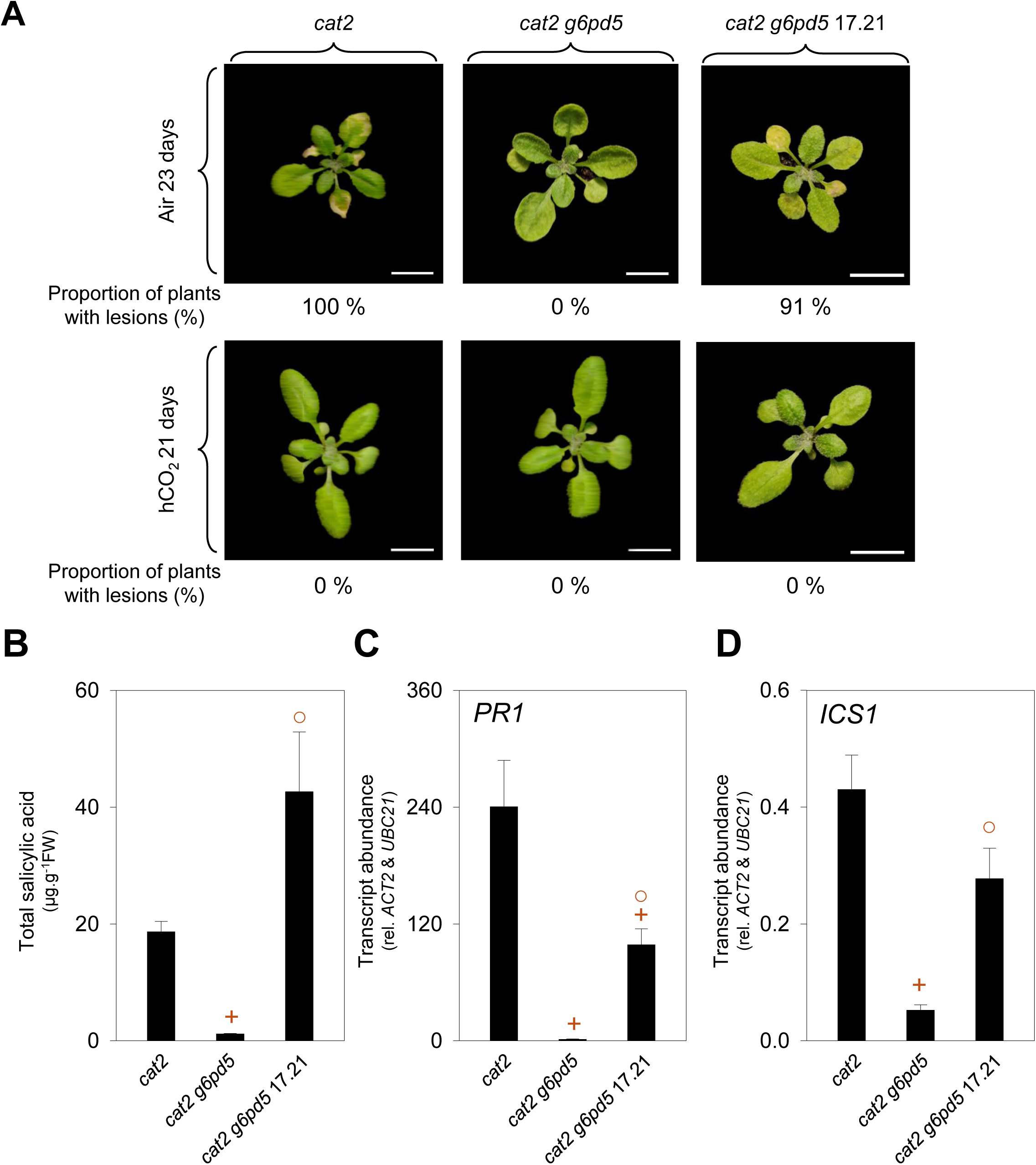
Characterization of the lesion phenotype and SA signaling in the *cat2 g6pd5* 17.21 revertant line. A. Representative photographs of the plants with lesions after 23 days of growth in air (400 ppm CO_2_) or 21 days of growth at high CO_2_ (3000 ppm CO_2_). Numbers below each picture indicate the percentage of plants presenting lesions (at least 30 plants in each case). Images are presented at different magnifications to clearly display lesions morphology. Scale bars correspond to 1 cm. B. Quantification of the total salicylic acid (SA) in the three genotypes (*cat2*, *cat2 g6pd5* and *cat2 g6pd5* 17.21). C & D. Quantification of *PR1* and *ICS1* transcript abundance by RT-qPCR. Transcript abundance was normalized to *ACT2* and *UBC21*. Analyses were performed on leaf extracts of 21-day-old plants grown in air and long day unless stated differently. Data are means ± SE of three biological replicates, except for SA contents, for which four replicates were used. Plus signs (+) indicate significant differences relative to *cat2*, and open circles (○) indicate significant differences relative to *cat2 g6pd5* (Student’s *t*-test, *p* < 0.05).

**Table 1.** Summary of *cat2 g6pd5* revertant lines with their CO_2_ dependence Summary table of the proportion of plants with lesions for each line according to growing conditions (air, 400 ppm CO_2_; high CO_2_, 3000 ppm CO_2_) and candidate genes identified by the sequencing. The proportion of plants with lesions according to growing conditions is shown in percentages.

| Line | Proportion of plant with lesions (%) |  | Candidate causal genes (if known) |  |
| --- | --- | --- | --- | --- |
|  | Air | High CO<sub>2</sub> | AGI | Gene name |
| 9.11 | 95 | 100 | *AT4G39800* | *MIPS1* |
| 13.27 | 100 | 100 | *AT4G39800* | *MIPS1* |
| 17.21 | 91 | 0 | *AT3G02360* | *PGD2* |
| 19.17 | 96 | 88 | *AT5G64930* or *AT5G61520* | *CPR5* or *STP3* |
| 21.14 | 100 | 100 | *AT4G39800* | *MIPS1* |

In *cat2 g6pd5 17.21*, in which lesion formation was observed in air but not hCO_2_, further analysis revealed that the lesion phenotype in air was accompanied by increased *PR1* and *ICS1* transcript abundance and elevated SA levels compared to *cat2 g6pd5* (Fig. 3B-D). These data demonstrate restoration of SA-associated responses in this revertant line. Sequencing data were consequently further analyzed to identify the causal mutation. This identified a single nucleotide polymorphism (SNP) in *PGD2*, encoding the cytosolic 6PGDH, the third enzyme of the OPPP in Arabidopsis. These results provide a first indication that PGD2 is important in oxidative stress-related signaling. The SNP introduces a non-synonymous substitution (E97K) within the predicted NADP(H)-binding domain (Fig. 4A) (Blum et al., 2025). To evaluate the potential structural impact of this substitution, a three-dimensional model of the PGD2 homodimer was generated using AlphaFold Multimer and aligned with the experimentally resolved structure of *Lactococcus lactis* GND (PDB: 2iyp) (Sundaramoorthy et al., 2007). The generated 3D structure indicates that E97 participates in folding and contributes to the proximity of the β-sheets of the NADP(H)-binding pocket, likely forming ionic interactions with the surrounding amino acids K71, R73 and D100 (Fig. 4B, Supplementary Figure S3). The E97K substitution might thus alter the local environment within the NADP(H)-binding pocket and potentially impair the enzyme activity.

**Figure 4.**
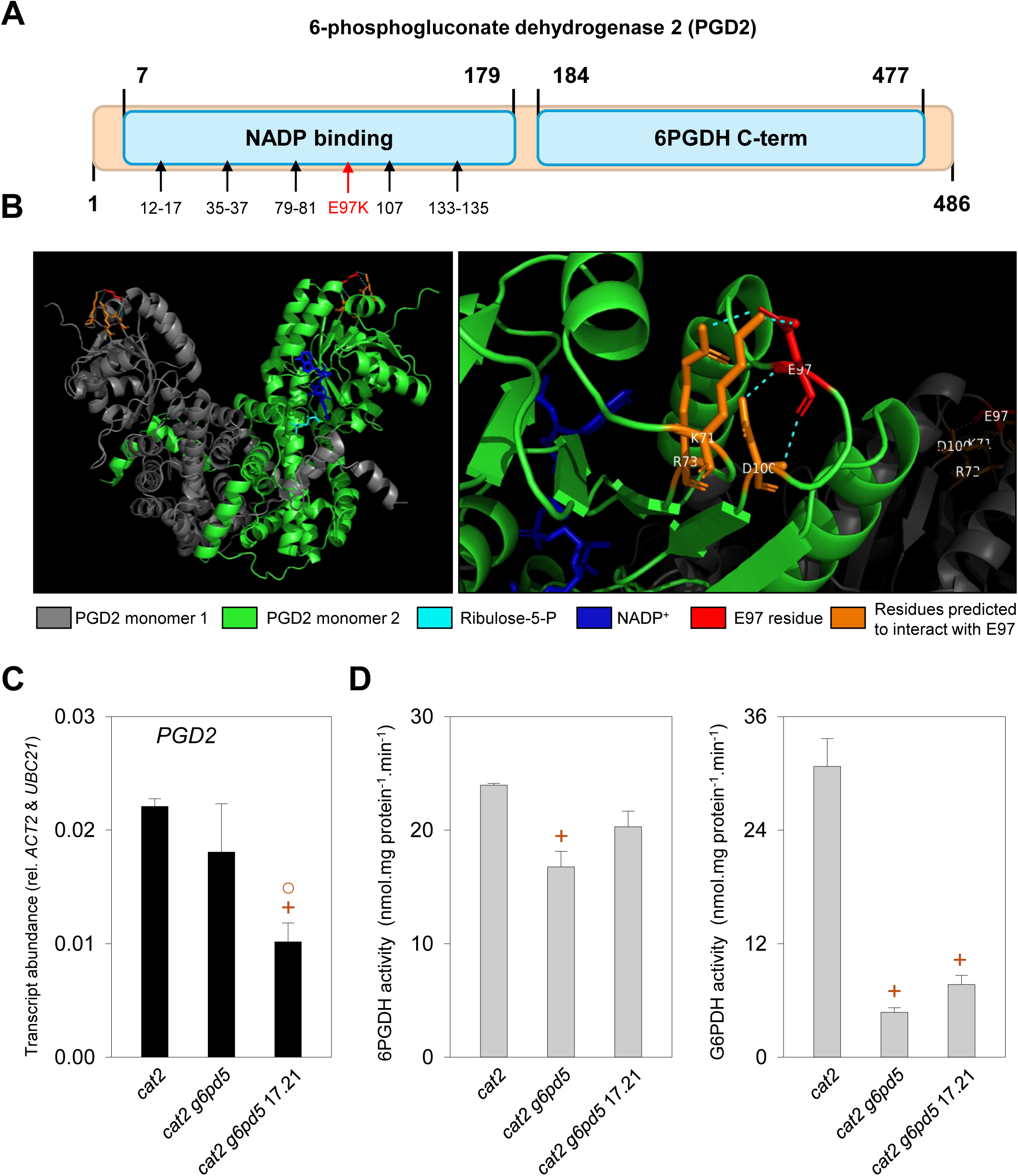
The impact of *cat2 g6pd5* 17.21 revertant line SNP on PGD2. A. Predicted Pfam domains and PIRSF key residues of the NADP(H) binding domain according to InterPro for PGD2 (AT3G02360; Q9FWA3) (www.ebi.ac.uk/interpro/). The mutated residue and corresponding substitution, E97K, is indicated in red. B. 3D mapping of E97 (highlighted in red) in the structure of the PGD2 homodimer predicted using AlphaFold-Multimer v3 (monomers are in green or grey). Left: General structure of the homodimer; Right: Zoom on E97 localization and its importance for the 3D structure. The Alphafold-Multimer v3 model was generated from the protein sequence employing MMseqs2 searches against UniRef plus environmental databases, utilizing both paired and unpaired MSAs, and executing six rounds of recycling. The structure was then aligned with the experimentally solved ortholog from PDB entry 2IYP to position the NADP_+_ cofactor (colored blue) and the reaction product Ru5P (colored cyan). Foldseek was used to identify the best ortholog with a structure that includes ligands in the PDB. Surrounding amino acids K71, R73, and D100 (colored orange) were identified based on their 3D spatial proximity to E97. Cyan dashed lines illustrate close interatomic distances, indicative of possible hydrogen bonds or contacts, as determined in the PyMOL visualization. C. Quantification of *PGD2* transcript abundance. Transcript abundance was normalized to *ACT2* and *UBC21*. D. Extractable activities of 6PGDH and G6PDH. Analyses were performed on leaf extracts of 21-day-old plants grown in air and long day for *cat2*, *cat2 g6pd5* and *cat2 g6pd5* 17.21. Data are means ± SE of three biological replicates. Plus signs (+) indicate significant differences relative to *cat2*, and open circles (○) indicate significant differences relative to *cat2 g6pd5* (Student’s *t*-test, *p* < 0.05).

However, despite this predicted structural alteration, only a difference in *PGD2* transcript abundance was measured. No difference in the total leaf extractable activities of either 6PGDH or G6PDH was detected between *cat2 g6pd5 17.21* and *cat2 g6pd5* (Fig. 4C and D). It is possible that compensatory effects of the other 6PGDH isoforms might explain the similar extractable activities. In a first approach to confirm the importance of PGD2, we attempted to complement the *cat2 g6pd5 17.21* revertant with the wild-type *PGD2* sequence. Unfortunately, despite repeated attempts, it was not possible to recover viable transformants expressing the gene in this background. These observations prompted us to further analyze the contribution of PGD2 to redox signaling using independent genetic backgrounds and approaches.

### Functional analysis of an allelic mutant confirms the role of *PGD2* in oxidative stress responses

Complete loss-of-function alleles of *PGD2* have previously been reported to be lethal (Hölscher et al., 2016; Doering et al., 2024). Therefore, we examined lines carrying T-DNA insertions either in the 5’ untranslated region or near the STOP codon (Supplementary Figure S4A). Homozygous mutants were recovered only for the SALK_033401 line, hereafter named *pgd2-5* (insertion in the 5’ untranslated region). In this line, total extractable 6PGDH activity was decreased by 14% compared to Col-0, whereas G6PDH activity was not affected (Supplementary Figure S4B and C). This modest decrease in extractable 6PGDH activity did not impact the phenotype in plants grown in standard conditions in air.

To study whether the *pgd2-5* mutation affected oxidative stress responses, we crossed *pgd2-5* with *cat2* to generate the *cat2 pgd2-5* double mutant. Surprisingly, given the wild-type phenotype of the *pgd2-5* single mutant, the *cat2 pgd2-5* mutant proved to be difficult to obtain, with significant genetic effects already apparent in the F2 generation of the double mutant cross. The cross resulted in a deviation from Mendelian segregation in the F2 generation, with *cat2 cat2 pgd2-5 pgd2-5* double homozygotes exhibiting seedling lethality at the cotyledon stage (Table 2). In addition, although *cat2 cat2 PGD2 pgd2-5* plants could be recovered, they displayed more extensive leaf lesions and decreased growth compared to *cat2 cat2 PGD2 PGD2* (Table 2). These observations pointed to an exquisite sensitivity of plants to *PGD2* expression levels in the *cat2* oxidative stress background.

**Table 2.** Segregation study of F2 for the *cat2* × *pgd2-5* cross Segregation study of F2 for *cat2* × *pgd2-5* grown in air in long days at a light intensity of 200 µmol photon/m²/s. A Monte Carlo exact multinomial test was performed for a predicted ratio of 1:2:2:1:1:4:2:2:1 for the genotypes in descending order in the table (*p*-value threshold 0.05). The phenotype of each genotype after 23 days of growth is described in the table.

| Genotype | Number of plants | Proportion (%) | Theoretical number of plants | Theoretical proportion (%) | Phenotype after 23 days of growth |
| --- | --- | --- | --- | --- | --- |
| CAT2 CAT2 PGD2 PGD2 | 19 | 10.38 % | 11 | 6.25 % | Wild-type phenotype |
| CAT2 *cat2* PGD2 PGD2 | 20 | 10.93 % | 23 | 12.5 % | Wild-type phenotype |
| CAT2 CAT2 PGD2 *pgd2-5* | 19 | 10.38 % | 23 | 12.5 % | Wild-type phenotype |
| *cat2 cat2* PGD2 PGD2 | 10 | 5.46 % | 11 | 6.25 % | 80 % of the plants with lesions |
| CAT2 CAT2 *pgd2-5 pgd2-5* | 16 | 8.74 % | 11 | 6.25 % | Wild-type phenotype |
| CAT2 *cat2* PGD2 *pgd2-5* | 66 | 36.07 % | 46 | 25 % | Wild-type phenotype |
| CAT2 *cat2 pgd2-5 pgd2-5* | 15 | 8.2 % | 23 | 12.5 % | Wild-type phenotype |
| *cat2 cat2* PGD2 *pgd2-5* | 11 | 6.01 % | 23 | 12.5 % | Growth delay and lesions observed on all plants |
| *cat2 cat2 pgd2-5 pgd2-5* | 7 | 3.83 % | 11 | 6.25 % | All plants dead at the cotyledon stage |
| Total | 183 | 100 | 183 | 100 |  |
| p-value | 0.00063 | < | 0.05 | Non-Mendelian segregation |  |

To determine whether the strong effect on growth of *pgd2-5* in the *cat2* background was linked to oxidative stress and to attempt to obtain viable double homozygous mutants, F2 progeny from the *cat2*×*pgd2-5* cross were grown in hCO_2_ to suppress oxidative stress linked to the *cat2* mutation (Fig. 5A). Genotyping of F2 plants grown at hCO_2_ confirmed that, in contrast to plants grown in air, viable *cat2 pgd2-5* double homozygotes could be readily recovered in this condition (Fig. 5B). To further assess the dependence of the phenotype on oxidative stress, plants were either grown under hCO_2_ for 26 days or grown for 18 days under hCO_2_ and then transferred to air for 8 days (Fig. 5A). No lesion phenotype was observed for any genotype when plants were grown continuously under hCO_2_. In contrast, transfer to air triggered lesion formation in *cat2* and *cat2 pgd2-5* (Fig. 5C). Notably, all *cat2 pgd2-5* plants displayed extensive lesions, whereas only 40% of *cat2* plants showed any lesions at the same time point (Fig. 5C). The extreme phenotype of *cat2 pgd2-5* was associated with smaller rosettes eight days after onset of oxidative stress, together with a higher proportion of leaf area exhibiting lesions compared to *cat2* (Fig. 6A). These effects were positively correlated with over-accumulation of *PR1* transcripts and higher SA accumulation in *cat2 pgd2-5* (Fig. 6B and C), corroborating the evidence from the screen that *PGD2* contributes to SA signaling in the context of oxidative stress.

**Figure 5.**
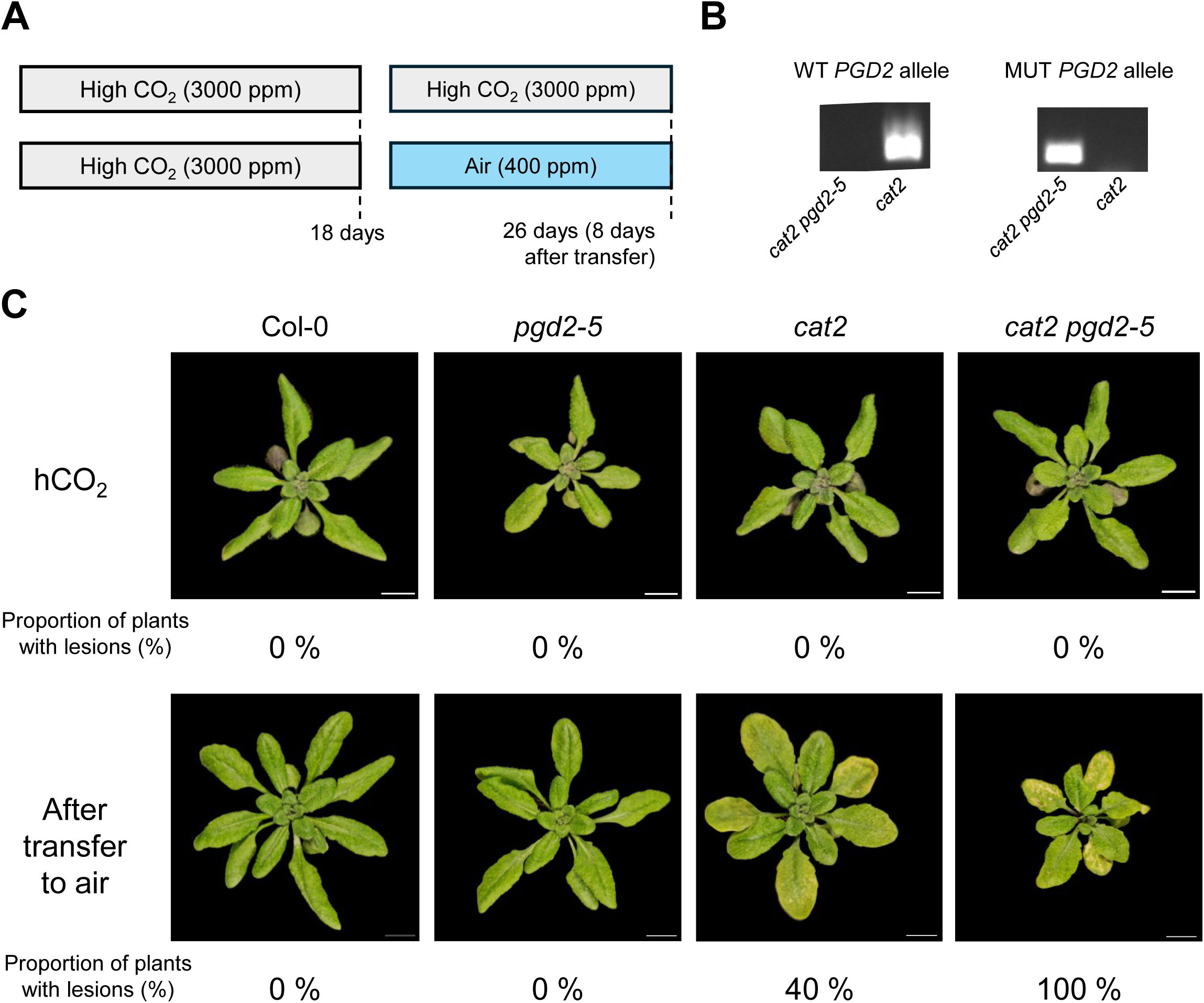
High CO_2_-dependent generation of the *cat2 pgd2-5* double homozygotes. A. Scheme depicting the experimental design used to identify and characterize the *cat2 pgd2-5* homozygote mutants. Plants were grown under ambient air (400 ppm) or high CO_2_ (3000 ppm) conditions, the latter suppressing photorespiratory H₂O₂ production. Plants were grown in long-day conditions with a light intensity of 200 µmol.m_-2_.s_-1_. The number of days corresponds to the days after the transfer of the seeds to growing conditions. B. Genotyping example of *cat2* and *cat2 pgd2-5* for the *PGD2* gene. Ethidium bromide signal of electrophoresis for PCR product of primers targeting the *PGD2* wild-type allele (WT *PGD2* allele) and the *pgd2-5* mutant allele (MUT *PGD2* allele). C. Representative phenotypes 8 days after transfer to air for Col-0, *pgd2-5*, *cat2* and *cat2 pgd2-5* (Control plants were kept at hCO_2_). Scale bars correspond to 1 cm. The proportion of plants with lesions in percentage at 8 days after transfer is indicated below the corresponding photograph for each genotype.

**Figure 6.**
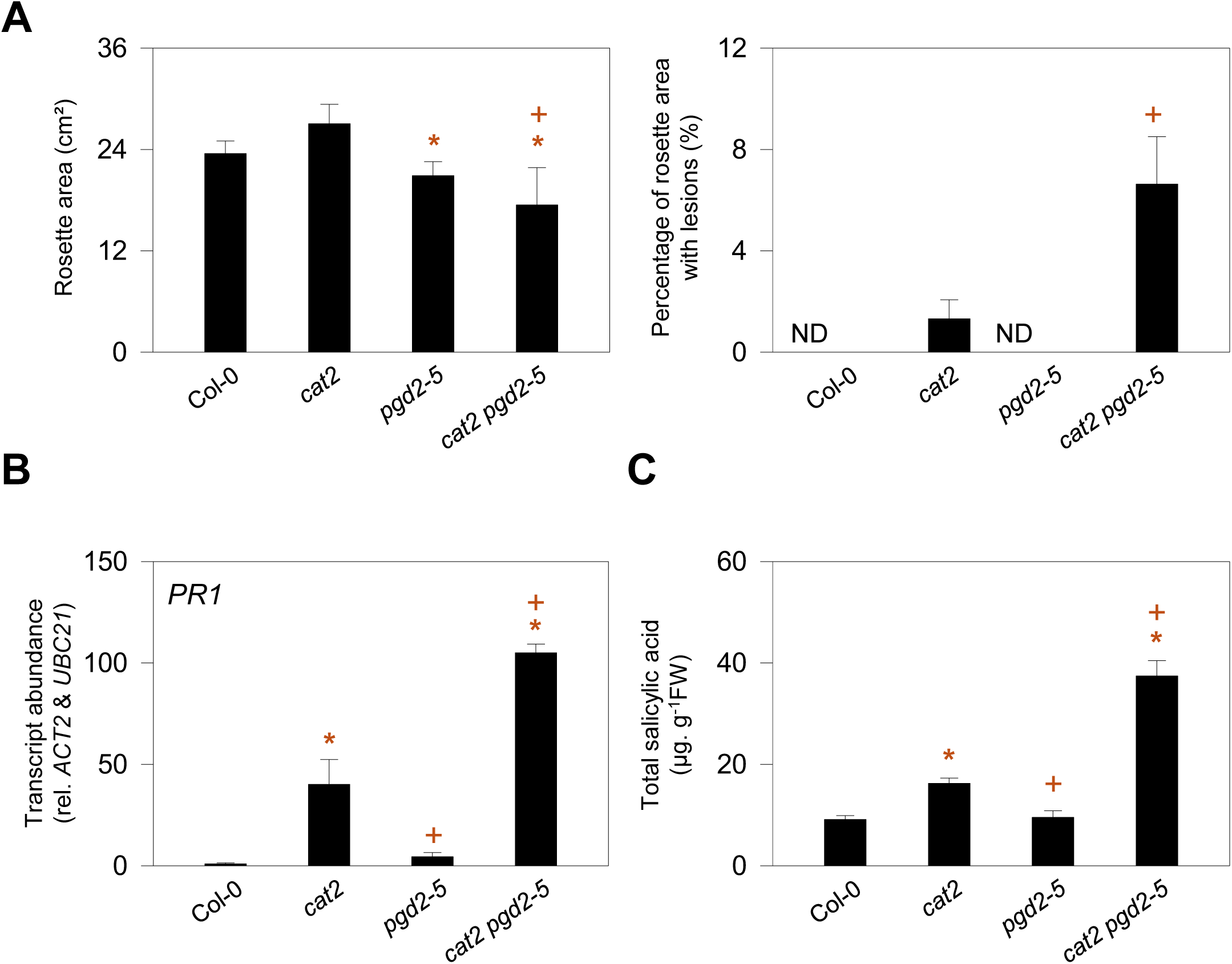
Analysis of lesion phenotype and SA-related pathogenesis responses in *cat2 pgd2-5* plants. A. Rosette area and percentage of rosette area displaying lesions in the indicated genotypes. Five plants were analyzed per biological replicate (15 plants in total per genotype). ND. Not detected. B. Transcript abundance for the SA marker gene, *PR1*, measured by RT-qPCR and normalized to *ACT2* and *UBC21* reference genes. C. Total salicylic acid content in the four genotypes. Analyses were performed on leaf extracts of plants grown from seeds for 18 days in 3000 ppm CO_2_ (high CO_2_) and then 8 days in air (400 ppm CO_2_). Data are means ± SE of three biological replicates, except for SA, where four replicates were used. Asterisks (*) indicate significant differences relative to Col-0, and plus signs (+) indicate significant differences relative to *cat2* (Student’s *t*-test, *p* < 0.05).

### Overexpression of *PGD2* in the *cat2* background abolishes SA signaling

Given the sensitivity of *cat2* phenotypes to the *pgd2-5* mutation, we hypothesized that overexpression of *PGD2* in the *cat2* background would produce opposite effects. A construct encoding PGD2 fused to GFP at the C-terminus was introduced into *cat2* plants. Two independent homozygous *cat2* PGD2 T3 lines were selected and analyzed to verify the transgene expression. Compared to *cat2*, both lines showed strong expression of the transgene, as indicated by *GFP* transcript detection and enhanced *PGD2* transcript abundance (Fig. 7A-B). Consistent with the predicted localization, PGD2-GFP fluorescence showed a signal typical of a cytosolic distribution in confocal microscopy analysis (Fig. 7C).

**Figure 7.**
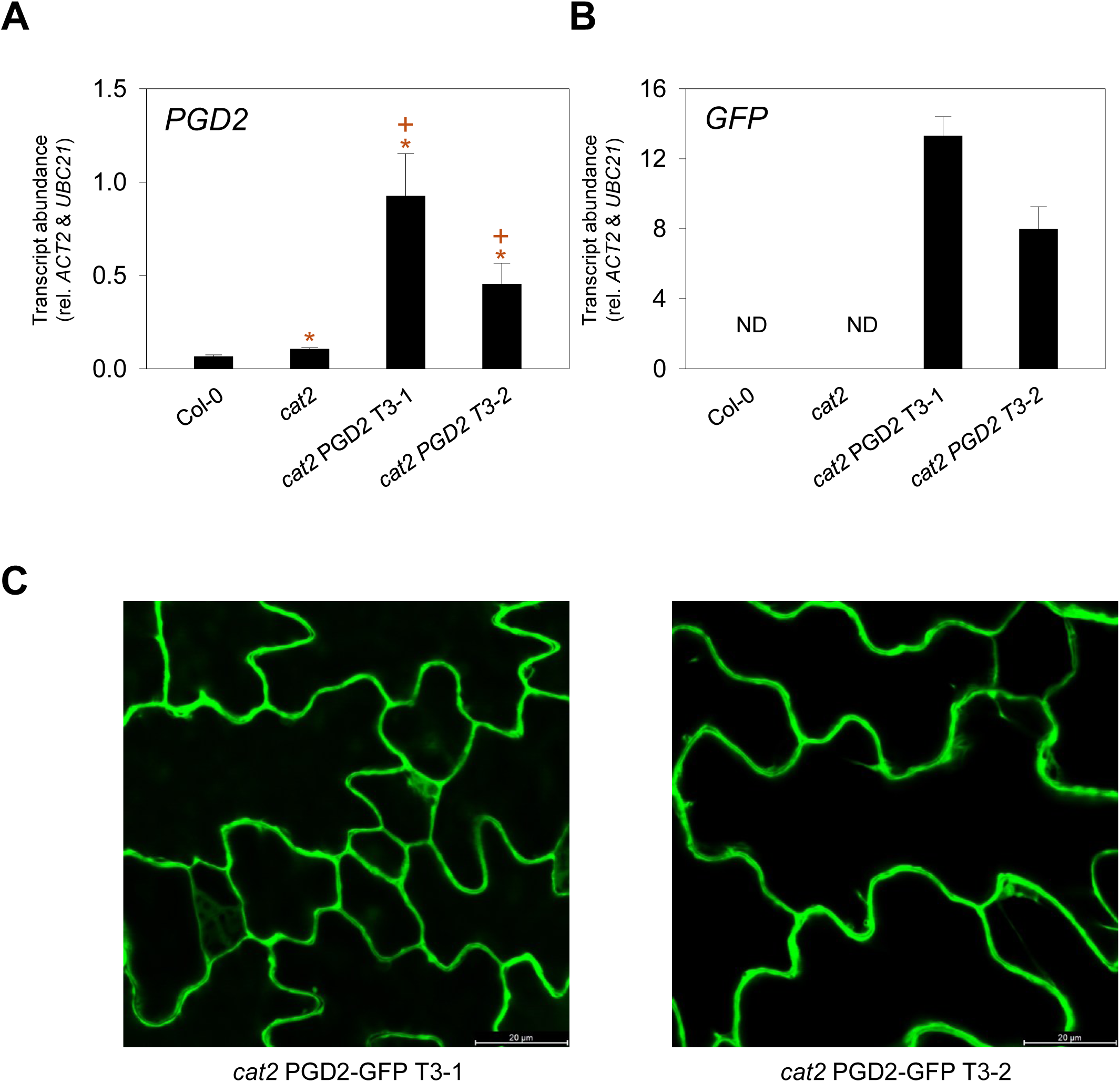
Generation and characterization of *cat2* lines overexpressing *PGD2*. A. Transcript abundance of *PGD2* in the indicated genotypes. B. Transcript abundance of GFP in the same lines. ND, not detected. Transcript levels were determined by RT-qPCR and normalized to the reference genes *ACT2* and *UBC21*. Analyses were performed on leaf extracts of 21-day-old plants grown in air and long day conditions. Data are means ± SE of three biological replicates. Asterisks (*) indicate significant differences relative to Col-0, and plus signs (+) indicate significant differences relative to *cat2* (Student’s *t*-test, *p* < 0.05). C. Representative confocal microscopy images showing GFP fluorescence in *cat2* PGD2-GFP lines.

In both transgenic lines, *PGD2* overexpression resulted in a more than four-fold increase in extractable 6PGDH activity (Fig. 8A) but only a minor increase in G6PDH activity (Fig. 8B). Because PGD2 is an NADPH-generating enzyme, we examined markers of cellular redox status in the overexpressors. Despite the increase in extractable activity, no significant differences were detected in oxidative stress markers: both abundance of *GSTU24* transcript and glutathione status were similar to *cat2* (Fig. 8C-D).

**Figure 8.**
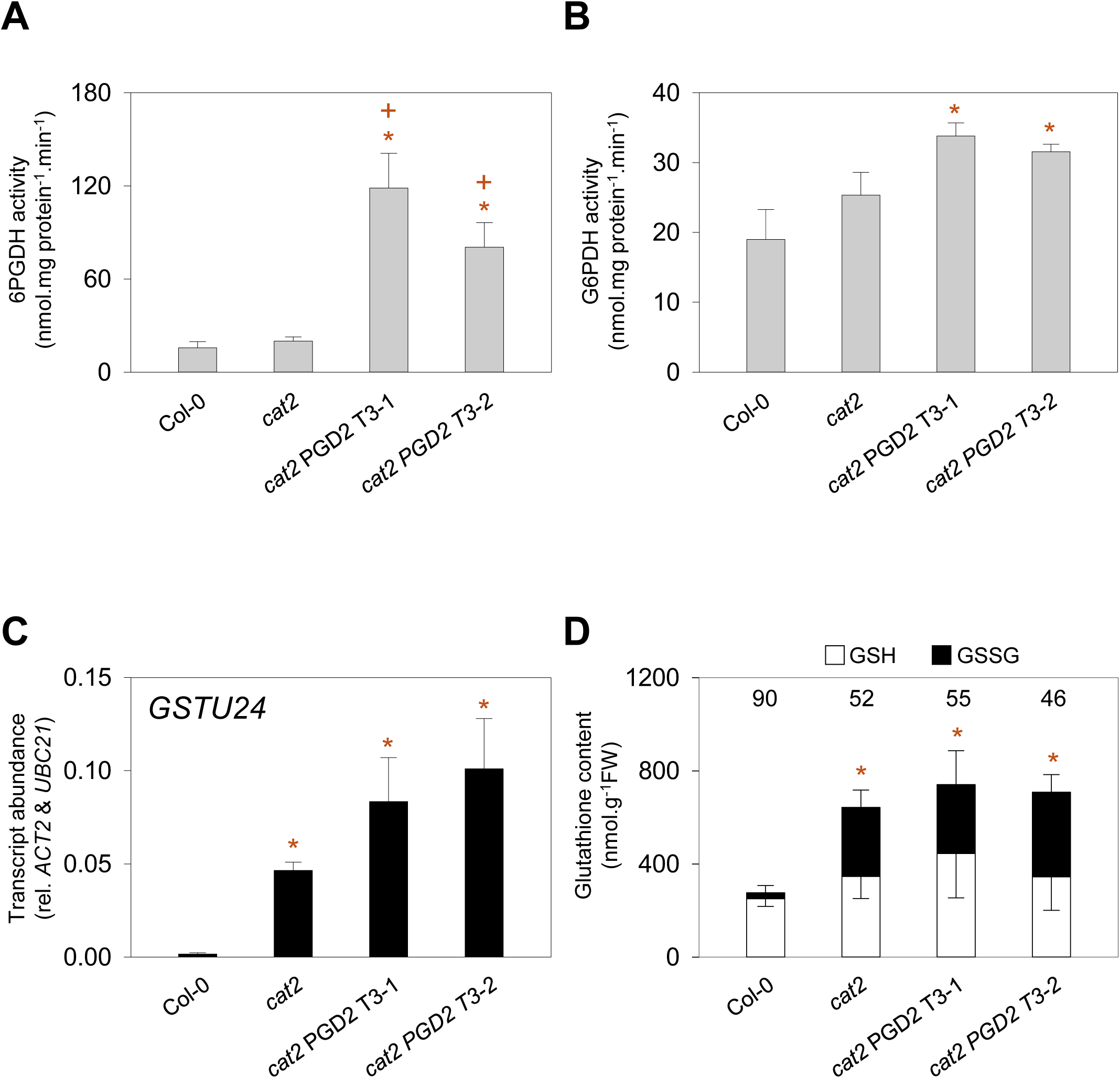
OPPP dehydrogenase activities and cellular redox state in *cat2* lines overexpressing *PGD2*. A. Extractable 6-phosphogluconate dehydrogenase (6PGDH) activity. B. Extractable glucose-6-phosphate dehydrogenase (G6PDH) activity. C. Transcript abundance of the oxidative stress marker gene *GSTU24*, determined by RT-qPCR. Transcript levels were normalized to the reference genes *ACT2* and *UBC21*. D. Glutathione content and redox state. Reduced glutathione (GSH) is shown as the white lower bars and oxidized glutathione (GSSG) as the black bars. The percentage of total glutathione present in the reduced form (% reduced) is indicated above each bar. Analyses were performed on leaf extracts of 21-day-old plants grown in air and long day conditions. Data are means ± SE of three biological replicates. Asterisks (*) indicate significant differences relative to Col-0, and plus signs (+) indicate significant differences relative to *cat2* (Student’s *t*-test, *p* < 0.05).

We next assessed the consequences of *PGD2* overexpression on SA-dependent responses. Strikingly, neither of the *cat2* PGD2 lines showed lesions (Fig. 9A), ie, the increased PGD2 activity produced the same effect as loss of *G6PD5* function in *cat2 g6pd5* (Trémulot et al. companion article). This suppression of lesions was accompanied by strongly decreased *PR1* and *ICS1* transcript levels, and decreased SA accumulation compared to *cat2* (Fig. 9B-D). These results confirm that PGD2 has effects opposite to those of G6PD5 on oxidative stress-induced SA signaling. Because both enzymes contribute to NADP_+_ reduction, the contrasting effects are difficult to explain simply by changes in NADPH availability. Instead, these findings suggest that another feature of OPPP metabolism contributes to the control of SA signaling during oxidative stress.

**Figure 9.**
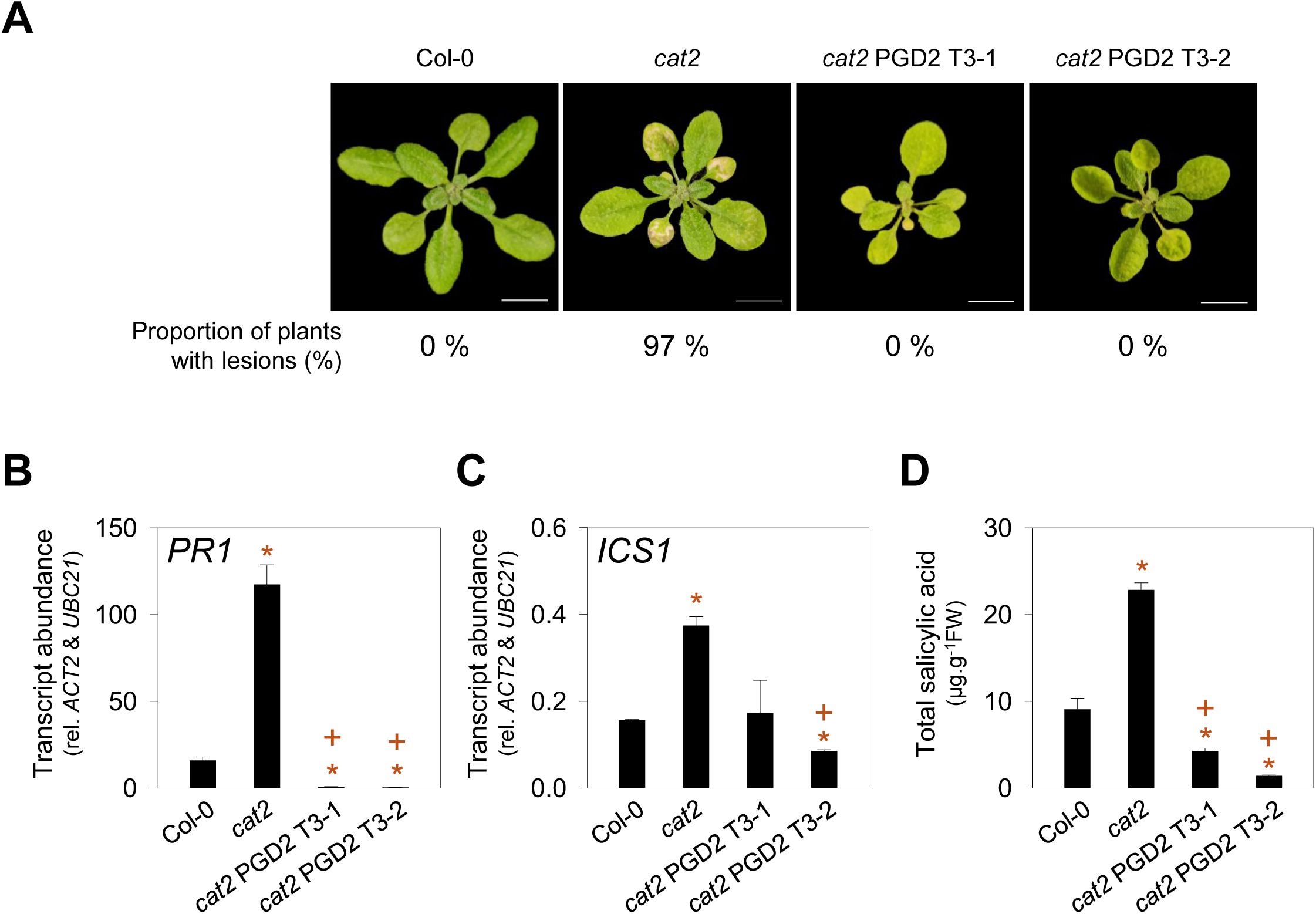
Characterization of lesion phenotype and SA-related responses in *cat2* lines overexpressing *PGD2*. A. Representative rosette phenotype of 21-day-old plants. The proportion of plants with lesions is indicated below the plant photograph (at least 34 plants per genotype). Scale bars correspond to 1 cm. B and C. *PR1* and *ICS1* transcript abundance. Transcript abundance was normalized to reference genes *ACT2* and *UBC21*. Data are means ± SE of three biological replicates. D. Total salicylic acid content. Analyses were performed on leaf extracts of 21-day-old plants grown in air and long day conditions. Asterisks (*) indicate significant differences relative to Col-0, and plus signs (+) indicate significant differences relative to *cat2* (Student’s *t*-test, *p* < 0.05).

### Evidence that an OPPP intermediate couples oxidative stress to SA signaling

In view of the antagonistic role of the two cytosolic dehydrogenases of the OPPP, G6PD5 and PGD2, we hypothesized that a metabolic intermediate produced by G6PDH upstream of 6PGDH might participate in SA signaling triggered by oxidative stress. Indeed, the accumulation of such a metabolite might be expected to be weakened in *cat2 g6pd5* and enhanced in *cat2 pgd2-5*. In the *cat2* PGD2 lines, more efficient metabolism might also antagonize its accumulation, hence producing a similar effect to that in *cat2 g6pd5*. Although technical challenges prevented accurate quantification of OPPP intermediates, GC-MS profiling has shown that oxidative stress in *cat2* causes dramatic accumulation of gluconate, an effect that is much less or not apparent in *cat2 g6pd5* or other *cat2* genotypes impaired in SA signaling (Chaouch et al., 2012; Han et al., 2013; Trémulot et al., companion article).

To test the possible influence of the OPPP intermediates, 6-phosphogluconolactone (6PGL) and 6-phosphogluconate (6PG), a chemical complementation strategy was used. While 6PG is readily available, 6PGL is unstable in solution (Miclet et al., 2001; Sadet et al., 2018). Hence, we attempted to reconstitute 6PGL production through treatment with a solution containing G6PDH, G6P, and NADP⁺. First, we examined whether this treatment could restore SA signaling in *cat2 g6pd5* leaves, using *PR1* expression as a marker. While neither G6P nor G6PDH alone was able to induce *PR1* expression, and NADP_+_ alone had only a modest effect, treatment with G6PDH, G6P, and NADP_+_ dramatically induced *PR1* transcripts to more than 50-fold higher levels than in the mock control (Fig. 10A). Alongside the increased *PR1* expression, the combined G6PDH+G6P+NADP⁺ treatment resulted in a five-fold increase in SA levels (Fig. 10B). These effects required an active enzyme, since a mixture containing heat-inactivated G6PDH produced effects comparable to NADP⁺ alone (Supplementary Figure S5A). They also seemed not to be caused directly by gluconate, since treatment with this compound failed to induce *PR1* expression in *cat2 g6pd5* (Supplementary Figure S5A).

**Figure 10.**
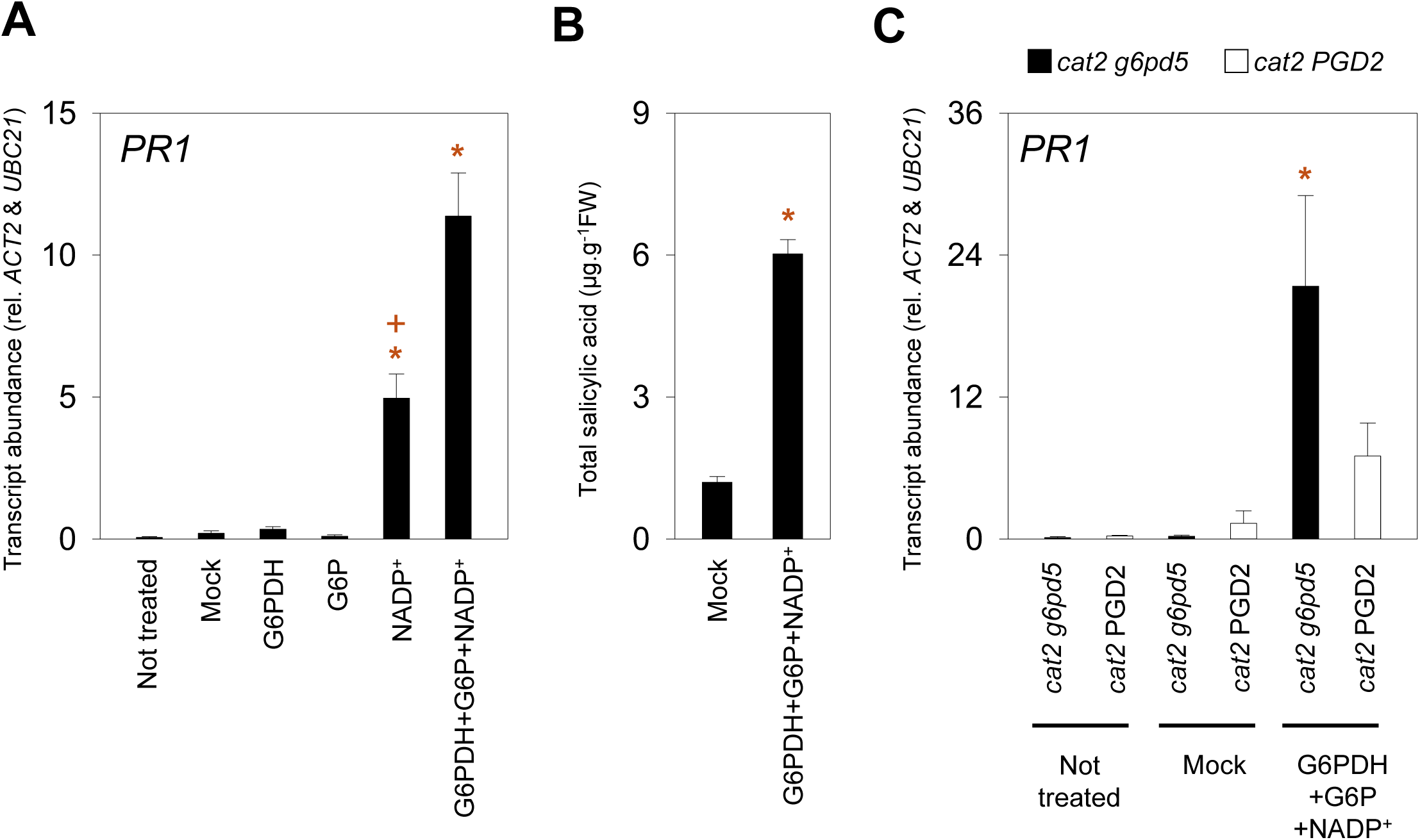
Effects of treatments stimulating the OPPP on SA signaling in *cat2 g6pd5* and *cat2* PGD2. A. Transcript abundance of the SA marker gene *PR1* measured 24 h after infiltration of the second leaf pair of *cat2 g6pd5*. B. Total SA content measured 24 h after infiltration of the second leaf pair of *cat2 g6pd5* plants. C. Transcript abundance of the SA marker gene *PR1* measured 24 h after infiltration of the second leaf pair of *cat2 g6pd5* (black) and *cat2* PGD2 (white). Sampled plants were grown for 22 days in air and long days at 200 µmol.m_-2_.s_-1_. Infiltration treatments contained yeast G6PDH (5 U/mL), glucose 6-phosphate G6P (15 mM), and NADP_+_ (600 µM). The *cat2* PGD2 corresponds to the line *cat2* PGD2 T3-1. Transcript abundance is normalized to *ACT2 and UBC21*. Data are means ± SE of three biological replicates. Asterisks (*) indicate significant differences relative to mock treatment, and plus signs (+) indicate significant differences relative to G6PDH+G6P+NADP_+_ treatments (Student’s *t*-test, *p* < 0.05).

We also assessed the effects of treating a *cat2* PGD2 overexpressor line, in which SA signaling is decreased in a similar manner to *cat2 g6pd5* but in which the strong increase in 6PGDH capacity (Fig. 8A) might be expected to antagonize accumulation of 6PGL and 6PG. The increase in *PR1* transcript abundance observed after the 6PGL-generating treatment (G6PDH+G6P+NADP⁺) in *cat2 g6pd5* was strongly attenuated in *cat2* PGD2 (Fig. 10C). Moreover, direct application of 6PG, which can be spontaneously formed from 6PGL and removed by 6PGDH (Miclet et al., 2001; Sadet et al., 2018), did not induce *PR1* expression (Supplementary Figure S5B). These results provide further evidence that metabolic processing within the OPPP modulates the strength of oxidative stress responses, and suggest that the signal is associated with 6PGL.

## Discussion

Oxidative stress is known to induce profound changes in cellular signaling, notably via pathways involving phytohormones. In the *cat2* mutant, in which oxidative stress occurs through increased availability of intracellular, metabolically produced H_2_O_2_, a key response is activation of SA accumulation and SA-dependent transcriptome reprogramming. The photorespiratory nature of the *cat2* mutant allows these responses to be abolished by high CO_2_. Several secondary mutations can also modulate the *cat2*-triggered SA response, even when oxidative stress is occurring in plants grown in air (Mhamdi et al., 2010a; Kerchev et al., 2016). Of these secondary mutations, we have identified only two that fully suppress SA signaling in our growth conditions. These are *sid2*, which abolishes SA biosynthesis (Chaouch et al., 2010), and *g6pd5*, reported here and in the companion manuscript. The present study shows that *PGD2* overexpression phenocopies these two loss-of-function mutations, and that this gene is important in linking intracellular oxidative stress to activation of SA signaling.

### *PGD2* plays a key role in oxidative stress-driven SA signaling

The genetic screen performed in the *cat2 g6pd5* background identified *PGD2* as a previously unrecognized component of the signaling pathway linking oxidative stress to activation of the SA pathway. The role of *PGD2* was confirmed through both loss- and gain-of-function approaches. In particular, overexpression of *PGD2* in the *cat2* background abolished SA accumulation and the associated lesion phenotype, whereas *pgd2* mutant alleles enhanced lesion formation and SA signaling (Fig. 3, 6 and 9). These opposing effects strongly support a regulatory role of PGD2 in oxidative stress signaling pathways. Complete loss of *PGD2* function has been reported to be lethal (Hölscher et al., 2016; Doering et al., 2024), suggesting that like the *pgd2-5* T-DNA allele, the *17.21* point mutation in the PGD2 sequence causes a partial loss of function. These viable alleles therefore provide useful genetic tools to investigate *PGD2* function in development or stress contexts where null mutants cannot be recovered.

Confocal analysis of PGD2-GFP showed that fluorescence signal accumulates in the cytosol (Fig. 7C). This is consistent with previous studies showing that C-terminal GFP fusions prevent the secondary targeting of PGD2 to the peroxisomes (Hölscher et al., 2016; Doering et al., 2024). Interestingly, the observation that cytosolic PGD2 overexpression is sufficient to suppress SA signaling in *cat2* indicates that its regulatory function operates primarily within this compartment. Thus, any secondary function or activity in the peroxisome seems to be of limited relevance in the context studied here.

### The two dehydrogenases of the cytosolic OPPP have an antagonistic role in oxidative stress responses

The present work establishes that the two enzymes acting sequentially within the cytosolic OPPP have antagonistic influences on SA signaling, revealing an unexpected level of regulatory complexity within the pathway. The striking contrast in the effects of the two enzymes on oxidative stress signaling is illustrated by the distinct effects of their overexpression in *cat2* contexts: while increased expression of the first enzyme promotes activation of the SA signaling, downstream *PGD2* expression opposes it (Fig. 8 and 9; Trémulot et al., companion manuscript).

A few previous studies have linked enzymes of the oxidative and non-oxidative branches of the PPP to SA signaling. The *pgl3-1* knockdown mutant, affecting the chloroplast-located OPPP, shows constitutive activation of SA responses (Xiong et al., 2009a; Lansing et al., 2020), whereas mutations affecting the cytosolic ribose-5-phosphate isomerase, *RPI2*, can trigger lesion phenotypes resembling the hypersensitive response (Xiong et al., 2009b). The regulation of the outcomes of oxidative stress by cytosolic OPPP enzymes echoes previous studies showing that cytosolic systems strongly influence the *cat2* phenotype. Although none of them impact *cat2*-triggered SA signaling as strongly as the effects reported here, loss of cytosolic *APX1*, *MDAR2*, *DHAR1/DHAR2* or *GR1* functions also have some effect.

### The role of the OPPP in SA signaling cannot be explained solely by NADPH production

Based on current concepts, a contribution of the OPPP enzymes to oxidative stress signaling would likely involve maintaining cellular redox homeostasis through NADPH production. For example, G6PDH and 6PGDH could generate NADPH necessary to sustain GR and/or MDHAR activity, thereby influencing the glutathione redox state, which in turn influences oxidative signaling (Mhamdi et al., 2010a; Xu et al., 2025).

While our analyses of glutathione and other factors show that G6PD5 can contribute to redox homeostasis (Trémulot et al., companion manuscript), such mechanisms are not sufficient to explain the impact of the OPPP on oxidative stress signaling. First, concerning glutathione redox state, our previous studies have shown that there is no simple relationship between this factor and SA signaling in the *cat2* background (Xu et al., 2025 and references cited therein). Second, comparative analyses of phenotypes and transcriptomes provide little evidence that loss of NADPH oxidase or ascorbate-glutathione pathway functions modulate *cat2*-triggered responses in the same way as *g6pd5* (Fig. 1). Third, both G6PD5 and PGD2 generate NADPH, yet the two enzymes modulate oxidative stress signaling in opposite ways. In the accompanying study, *cat2 g6pd5* mutants over-accumulate oxidized glutathione yet fail to activate SA signaling. Conversely, in the present work, *PGD2* overexpression suppresses SA signaling without substantially altering glutathione redox status. This latter observation could possibly reflect a strong limiting role of G6PD5 over operation of the pathway and, therefore, 6PGDH. In any case, the data provide little evidence that NADPH availability alone explains the impact of the OPPP on SA responses, and suggest that other cytosolic components or metabolic intermediates are involved.

### Evidence for a signaling role of an OPPP intermediate

If NADPH production alone cannot explain OPPP-mediated signaling during oxidative stress, an alternative possibility is that a metabolite of the pathway acts directly as a signaling molecule. The opposing phenotypes of *cat2 g6pd5* and *cat2 pgd2* lines are consistent with this idea. Because G6PD5 catalyzes the first oxidative step of the pathway whereas PGD2 acts downstream, the relevant signal is likely generated between the G6PDH and 6PGDH reactions, implicating intermediates such as 6PGL or 6PG.

Several experimental observations support the conclusion that a metabolite produced by G6PDH and removed by 6PGDH acts as a signaling intermediate. First, the *cat2 g6pd5 17.21* revertant line genetically links *G6PD5* and *PGD2* in the control of SA signaling. Second, unlike loss of *G6PD5* function, the *pgd2-5* mutation is lethal in the *cat2* background when photorespiration is active, indicating that accumulation of metabolites upstream of 6PGDH has major physiological consequences. In this connection, it should be noted that the *pgd2 17.21* allele, carrying a point mutation in the *PGD2* sequence, was identified in a G6PD5-deficient background. It is possible that without the attenuating effect of the *g6pd5* mutation on the cytosolic OPPP, this allele would be also lethal. Indeed, genetic interactions have been described in yeast and Drosophila that suggest strong interplay between the two enzymes in determining phenotypes: severe growth defects caused by loss of 6PGDH function can be suppressed by loss or manipulation of G6PDH activity (Gvozdev et al., 1977; Lobo and Maitra, 1982; Bertels et al., 2021; Bertels et al., 2025). Further evidence comes from pharmacological experiments, which point specifically towards 6PGL as a signaling molecule (Fig. 9; Supplementary Figure S5). Infiltration with a 6PGL-generating treatment (G6PDH + G6P + NADP_+_) restores SA signaling in *cat2 g6pd5*, whereas treatments with G6P, 6PG or gluconate alone have no effect. Importantly, this response is strongly attenuated in *cat2* PGD2 plants, possibly reflecting limited accumulation of 6PGL when 6PGDH capacity is enhanced.

It is noteworthy that 6PGL has interesting chemical features. The δ-6PGL directly generated by G6PDH rapidly equilibrates with a γ-6PGL isomer, which hydrolyses spontaneously to 6PG faster (Miclet et al., 2001; Sadet et al., 2018). Hydrolysis of δ-6PGL (but not γ-6PGL) is also catalyzed by 6-phosphogluconolactonase. Altered flux through the OPPP could favor transient accumulation of γ-6PGL, allowing this metabolite to function as a signal without strongly affecting downstream metabolism. Lactones are chemical groups that are known to be important in cellular signaling. Interestingly, γ-6PGL has recently been reported to act as a signaling molecule in human cells (Gao et al., 2019).

### A key role for metabolic flux in oxidative stress signaling?

One feature that could make γ-6PGL well suited to a signaling function is that, as a side-product of the pathway, its accumulation is only indirectly affected by G6PDH and PGL activities (Sadet et al., 2018). This could favor gradual changes in its concentration as a function of pathway flux and the relative control strength of the different enzymes. Increases in PPP cycling occur in the context of oxidative stress in animals, fungi, and bacteria (Christodoulou et al., 2018; Christodoulou et al., 2019; Hurbain et al., 2022; TeSlaa et al., 2023). In addition to any direct redox-signaling role that NADPH might play, its homeostasis is a key control point to consider in driving possible metabolite signaling. Indeed, NADPH inhibits both G6PDHs and 6PGDHs, and NADP_+_ binding to G6PDH was shown to favor its tetrameric structure and thus its enzymatic activity (Rosemeyer, 1987; Au et al., 2000; Wakao and Benning, 2005). Such regulation may therefore help to explain the effects of NADP_+_ treatment (Fig. 9, Supplementary Figure S5) (Zhang and Mou, 2009).

The OPPP is at the crossroads of carbon metabolism and redox homeostasis. If G6P concentrations or enzyme capacities are not strongly limiting, flux through the OPPP could be largely controlled by NADPH/NADP_+_ turnover. In this case, a major driver of flux through the pathway would be increased NADPH consumption. During oxidative stress, increased NADPH demand associated with both ROS production and antioxidant metabolism would favor lowered NADPH:NADP_+_ ratios, stimulating G6PDH activity and increasing overall flux through the pathway, possibly leading to accumulation of putative signaling molecules, such as γ-6PGL, which by binding to sensitive proteins could activate phytohormone signaling (Fig. 11). Restoration of redox homeostasis, including NADPH:NADP_+_ ratios, would cause gradual attenuation of the signal as flux decreases and the metabolite returns to basal levels. Our observations raise the possibility that, within the context of oxidative stress, the cytosolic OPPP acts not only as a metabolic source of NADPH but also as a signaling module linking redox metabolism to activation of immune pathways. Mechanisms such as these could provide links between ROS and signaling pathways that occur via indirect effects on metabolic flux, providing an additional input to those that involve direct redox modifications (Fig. 11).

**Figure 11.**
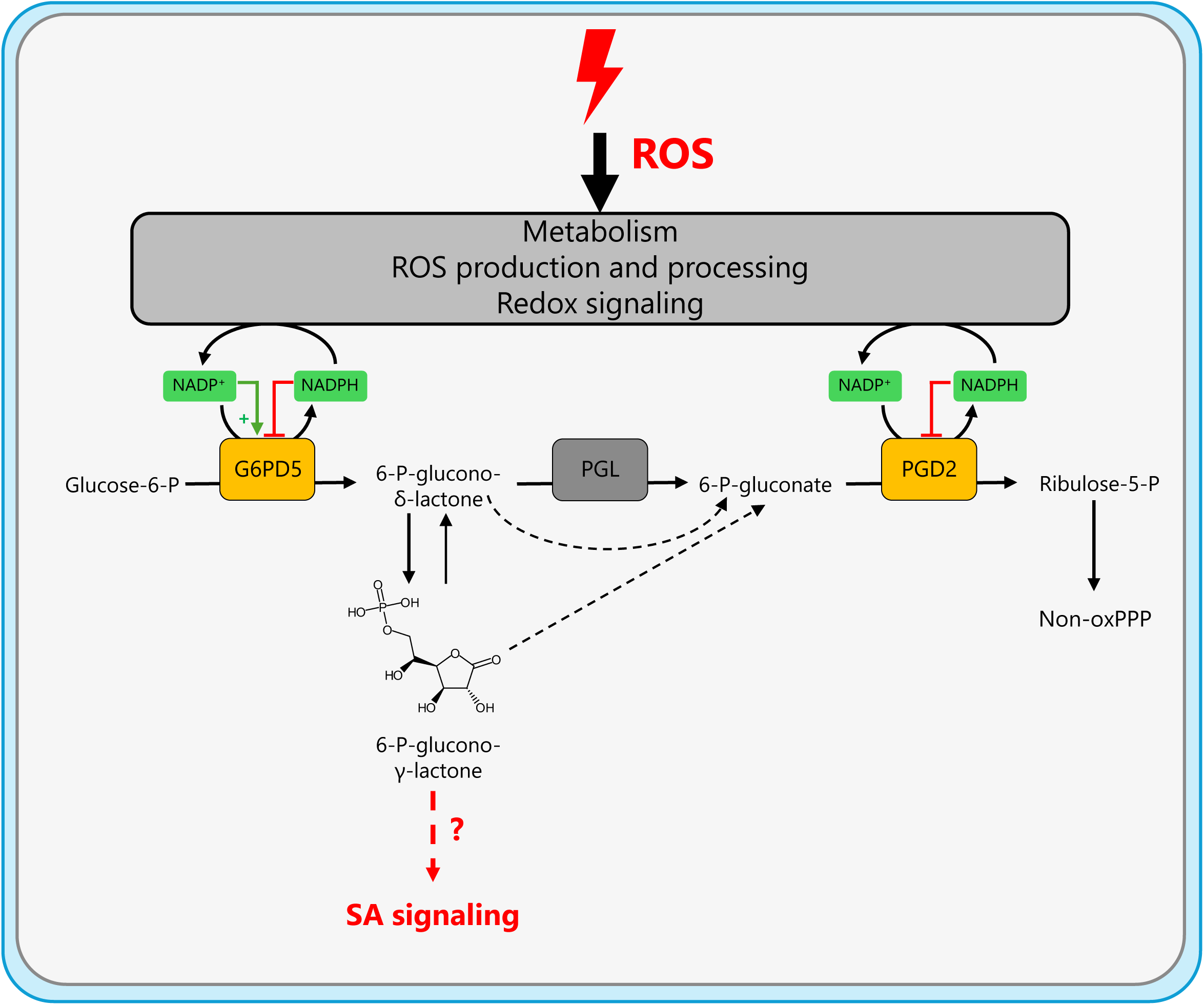
Model linking oxidative stress, cytosolic OPPP activity, and SA signaling. Enhanced NADP(H) turnover during ROS production and detoxification is proposed to stimulate cytosolic OPPP flux, leading to increased accumulation of 6-phosphogluconolactone (6PGL). Elevated levels of 6PGL, particularly the γ-6PGL form, may function as signaling intermediates through an unknown mechanism (?). Activation of this pathway leads to SA-dependent responses and the development of hypersensitive-response-like lesions. Non-oxPPP, non-oxidative pentose phosphate pathway.

## Materials and methods

### Plant materials, growth conditions, and sampling

The T-DNA lines were all in the Columbia ecotype and were obtained from the European Arabidopsis Stock Center (NASC, www.arabidopsis.info). Published mutants studied here were *cat2* (SALK_076998; (Queval et al., 2007), *cat2 sid2* (Chaouch et al., 2010), *cat2 atrbohd*, *cat2 atrbohf* (Chaouch et al., 2012), *cat2 mdar2* (Xu et al., 2025), *cat2 dhar123* (Rahantaniaina et al., 2017), *cat2 g6pd5* (Trémulot et al., companion article), and *pgd2-1* (Hölscher et al., 2016). Mutants for PGD2 were identified using the SIGnAL website (signal.salk.edu/cgi-bin/tdnaexpress; Alonso et al., 2003) and correspond to GK-692E09 (*pgd2-3*), SALK_057911 (*pgd2-4*), and SALK_033401 (*pgd2-5*) (Supplementary Figure S4A). Plants were genotyped by analyzing PCR products of specific DNA amplification using gel electrophoresis with primers listed in Supplementary Table S1.

The plants grown on soil were sown on Jiffy pellets and, after 2 days of stratification, grown in a controlled environment. The 7-day-old seedlings were then transferred to 5.5 cm round pots containing soil (Tref terreau P1, Jiffy France SARL). The standard growing conditions were 22 °C under light, 20 °C in the dark, and 65% humidity. Plants were maintained in long days (LD; 16h light/8 dark) with light intensities of 200 μmol photon/m²/s at leaf level. The hCO_2_ and air conditions refer to 3000 ppm CO_2_ and 400 ppm CO_2_, respectively. The plants were watered three times per week and, after transfer to pots, given nutrient solution twice weekly (Plant-prod 14-12-32, 280 g/L and Fertiligo 4.35 mL/L). Unless stated otherwise, leaves of 21-day-old plants were harvested in the middle of the photoperiod, and material was stored at −70 °C until analysis. All data corresponds to at least three biological replicates.

For *in vitro* experiments, growth was performed in 12×12 cm petri dishes containing autoclaved ½ MS and 0.8 % agar medium. The seeds were surface-sterilized with hypochlorite-ethanol (1:9, v/v) and ethanol 95 %. After stratification, petri dishes were placed at a light intensity of 70 μmol photon/m²/s in LD at temperatures of 20 °C (light period) and 18 °C (dark period) and a humidity of 65 %.

### Shoot phenotype study

The presence of HR-like lesions was determined by direct observation of the plants and lesions were quantified by image analysis. Rosette area and leaf lesion area were determined using Fiji (Schindelin et al., 2012). Briefly, the background was manually cropped on scaled images, which allowed the determination of the rosette area. The lesion area was determined using Fiji’s RGB thresholding and manual selection for refinement, if required. The percentage of rosette area with lesions was determined from the lesion area and rosette area data.

### Transcript quantification using RT-qPCR

RNA was extracted using the NucleoSpin RNA Plant and Fungi kit, which included the on-column rDNase step, according to the manufacturer’s protocol (Macherey-Nagel). Reverse transcription was performed with 1 μg RNA, 0.5 μg oligodT, and following the manufacturer’s instructions for ImProm-II™ RT (Promega). The cDNAs obtained were diluted 25 times. For qPCR, the LightCycler_®_ SYBR Green I Master mix (Roche) was used following the manufacturer’s guidelines. The qPCR and analysis were done as in Queval et al. (2007) for 45 cycles and using a LightCycler_®_ 480 II and corresponding software (Roche). Normalized transcript levels were determined using predetermined primer efficiencies and *ACT2* and *UBC21* as reference genes, with each biological replicate calculated as the mean of three technical replicates.

### Total salicylic acid quantification

The total salicylic acid (SA) amount was determined as described in Trémulot et al., companion article. Briefly, the liquid nitrogen-frozen samples were ground, and extraction was conducted using methanol. The sample was purified with diethyl ether partitioning and resuspended in HPLC solvents. Total SA content was determined by HPLC with fluorescence detection (Waters) using peak areas and calibration curves generated from standards.

### Measurement of G6PDH and 6PGDH extractable activity

The G6PDH and 6PGDH activities were measured according to Mhamdi et al. (2022). In brief, proteins were extracted using a liquid nitrogen-frozen mortar and pestle, desalted using a pre-equilibrated NAP-5 column (Cytiva), and activity was measured using A340 in a UV-compatible cuvette (BRAND_®_ GMBH) into a Cary60 spectrophotometer (Agilent). The same protocol was used for both enzymes, except for the identity of the substrate (G6P for G6PDH, 6PG for 6PGDH). The protein content in the extract was determined using the Bradford method (Sigma) (Bradford, 1976). For each biological replicate, the value was taken as the mean of three technical replicates.

### Measurement of glutathione levels and redox state

Glutathione was quantified following the protocol of Noctor and Mhamdi (2022). Leaf tissue was extracted in 0.2 M HCl using a liquid nitrogen-frozen mortar and pestle, followed by neutralization with 0.2 M NaOH. Total glutathione and oxidized glutathione (GSSG) were determined by monitoring the reduction of DTNB in the presence of NADPH and GR, resulting in an increase in absorbance at 412 nm proportional to glutathione concentration. Assays were performed in UV-compatible 96-well plates (Costar_®_) using a TECAN Infinite M200 Pro plate reader (Tecan Life Sciences). For each biological replicate, values represent the mean of three technical replicates

### Genetic screen procedure

Seeds of *cat2 g6pd5* were mutagenized using ethyl methanesulfonate (EMS). The mutagenesis efficiency was determined as in Martín et al. (2009). Screening for appearance of lesions in M2 was conducted under conditions that allow lesions in *cat2*: i.e., LD and light intensity of 200 μmol photon/m²/s. Of approximately 70 000 M2 plants, those showing lesions were checked for homozygosity of the *cat2* and *g6pd5* alleles. To confirm the lesion phenotype, M3 were grown then backcrossed with *cat2 g6pd5*, if the M3 plant showed lesions. The F1 and F2 of this backcross were characterized for lesion phenotype and its segregation. Sequencing of revertant lines was done on a bulk sampling of 50 F2 individuals presenting lesions and was compared to the reference genome *cat2 g6pd5*. SHOREmap was used to identify causal mutations (Sun and Schneeberger, 2015). The occurrence of the lesion phenotype in the F3 of the backcrossed lines was studied by growing them either in air for 23 days or in hCO_2_ for 21 days.

### 3D modelling, structure similarity, and multiple sequence alignment

The PGD2 homodimer structure was predicted with AlphaFold-Multimer v3 (ColabFold implementation; Jumper et al., 2021; Evans et al., 2022; Mirdita et al., 2022) and visualized in PyMOL (Open-source distribution; v3.1.0). Foldseek was used to identify a protein with a structure that includes ligands in the PDB and showing the highest 3D structure similarity to the PGD2 Alphafold-generated structure (search.foldseek.com/search) (van Kempen et al., 2024; Kim et al., 2025). Multiple alignment was performed on the peptide sequence of PGD2 and the identified protein using Clustal Omega (www.ebi.ac.uk/jdispatcher/msa/clustalo), and visualization was done with Jalview (www.jalview.org).

### Vector construction, generation of transformed lines, and subcellular localization

*PGD2* was expressed as a C-terminal GFP fusion driven by the CaMV 35S promoter. For cloning, total RNA was extracted with ReliaPrep™ RNA Miniprep Systems from leaf tissue (Col-0) to obtain cDNA by reverse transcription with qScript™ cDNA Supermix (Quantabio). The CDS of *PGD2* was amplified with the appropriate primers (Supplementary Table S1) with PCR using iProof™ High Fidelity PCR kit (Bio-Rad). The sequence was then cloned into the pDONR™221 vector with Gateway_®_ BP protocol following the manufacturer’s instructions (Invitrogen). The LR recombination reaction was performed according to the supplier’s instructions with pH7FWG2 (Karimi et al., 2002). The construct was propagated in *E. coli* DH5α grown at 37 °C on LB medium with the appropriate antibiotics. Plasmid extraction was done using GeneJET™ Plasmid Miniprep kit following the manufacturer’s instructions (Thermo Fisher Scientific). Colony PCR was performed with ALLin™ Red Taq Mastermix (highQu) by analyzing product length. Constructs were verified by Sanger sequencing. The *PGD2*-containing vector was introduced into *A. tumefaciens* GV3101, and T0 plants were transformed using the floral dip method (Clough and Bent, 1998). T1 seedlings were selected *in vitro* on 30 µg/mL Hygromycin B and transferred to soil for seed production. T2 and T3 generations were similarly selected, and lines exhibiting resistance similar to that of single-locus insertion were selected. Expression of the transgene was measured in candidate T2 and T3 transformants by RT-qPCR on the *GFP* sequence, as well as the *PGD2* transcripts. Two independent lines were selected for further characterization. Subcellular localization was assessed by confocal microscopy of the GFP signal as described in Rahantaniaina et al. (2017), using a Leica Stellaris confocal microscope (Leica Microsystems).

### Pharmacological treatment experiment

Experiments involving pharmacological treatments were conducted on 21-day-old plants by infiltrating the indicated solutions into the abaxial surface of the second leaf pair using a needleless syringe. Treatments were performed in the middle of the photoperiod with a minimum of infiltration events until the entire leaf limb was visibly infiltrated. All solutions were prepared freshly and kept on ice until infiltration. The mock corresponds to the buffer mix present in all treatment solutions: 1 mM pH 7.4 HEPES, 5 mM NaCl, 2.5 mM MgCl2. G6PDH from yeast (Sigma) enzyme concentration was adjusted to 5 U/mL within the treatment solution. For enzyme deactivation (BoiledG6PDH), the prepared G6PDH was separated into two tubes, and the “BoiledG6PDH” was heat-denatured by a 15-minute incubation at 95 °C, followed by cooling on ice. The deactivation was verified by measuring G6PDH activity as described above. All other compounds (Sigma) were dissolved in water to final concentrations in the treatment solution of 15 mM G6P, 600 μM NADP_+_, 15 mM 6PG, and 15 mM gluconate. 24 h after the treatment, the treated or untreated second leaf pairs were sampled and stored at −70 °C until analysis.

### Transcriptomic analysis

Two transcriptomic experiments were analyzed. The data were obtained with microarray (experiment 1; Trémulot et al., companion article) and RNA-seq (experiment 2; Xu et al., 2025) techniques. For experiment 1 (microarray) analysis, total leaf RNA was extracted and hybridized to ATH1-121501 GeneChip_®_ arrays by the VIB Nucleomics Core (nucleomicscore.sites.vib.be) according to the manufacturer’s instructions (Affymetrix). Raw image files were analyzed following the maEndToEnd workflow (Klaus and Reisenauer, 2018). Quality control was performed using the arrayQualityMetrics 3.64.0 package (Kauffmann et al., 2009). Data preprocessing used the oligo 1.71.7 package (Carvalho and Irizarry, 2010). Low-intensity probes were filtered using a median log2intensity ≥ 4.5 in at least three samples. Annotations were retrieved from ath1121501.db using the AnnotationDbi 1.70.0. Differential expression analysis was performed with the limma 3.64.3 package by fitting linear models (lmFit) and applying empirical Bayes moderation (eBayes) (Ritchie et al., 2015). DEGs for a pairwise comparison were considered to be those with FDR < 0.01 and |log2FC| > 1 (Supplementary Table S2). For experiment 2 (RNA-seq), RNA was extracted from leaf tissue using the Spectrum Plant Total RNA Kit for each sample, including a DNase step (Sigma). The Nucleomics Core (VIB, Leuven, Belgium) performed library preparation with the TruSeq Stranded mRNA Library Preparation Kit (Illumina) and sequencing using Illumina NextSeq 500 (75-bp single-end reads). Read alignment on the *A. thaliana* genome was performed with STAR 2.5.2b (Dobin et al., 2013) using the Araport11 annotation (Cheng et al., 2017). The counts were quantified using Subread 1.6.2. (Liao et al., 2014). Genes with fewer than 5 reads in at least three biological replicates were excluded. DiCoExpress was used on counts for the quality control and determination of DEGs (Lambert et al., 2020). A DEG corresponds to FDR < 0.01 and |log2FC| > 1 (Supplementary Table S3). Heatmap construction was performed with the ComplexHeatmap 2.24.1 package (Gu, 2022), and Venn diagrams were generated using the eulerr 7.0.2 package.

### Statistical analysis and bioinformatics

All statistical bioinformatics analyses and graph generation were performed using Excel (Microsoft) and R 4.5.1 in RStudio. Monte Carlo exact multinomial test with a *p*-value of 0.05 was performed with the XNomial v1.0.4.1 R package. For annotation, Thalemine was used (bar.utoronto.ca/thalemine/), together with Uniprot (uniprot.org) and Interpro (www.ebi.ac.uk/interpro/). Reference sequences were retrieved from Araport11 using Phytozome (https://phytozome-next.jgi.doe.gov/). Protein tertiary structures were obtained from PDB (www.wwpdb.org; www.rcsb.org).

## Accession numbers

Genes specifically studied here are *CAT2* (*AT4G35090*), *G6PD5* (*AT3G27300*), *AtRBOHD* (*AT5G47910*), *AtRBOHF* (*AT1G64060*), *MDAR2* (*AT5G03630*), *DHAR1* (*AT1G19570*), *DHAR2* (*AT1G75270*), *DHAR3* (*AT5G16710*), *MIPS1* (*AT4G39800*), *CPR5* (*AT5G64930*), *STP3* (*AT5G61520*), *PGD2* (*AT3G02360*), *ICS1* (*AT1G74710*), *PR1* (*AT2G14610*), and *GSTU24* (*AT1G17170*).

## Acknowledgments and Funding

The authors thank Aleksandra Lewandowska for help with SHOREmap analysis.

## Author Contributions

G.N. conceived the original research project and supervised the experimental work with contributions from A.M. L.T. performed most of the experimental work and contributed hypotheses. Z.Y. and G.C-I. performed the genetic screen. E.I-B. contributed to data processing and interpretation. K.V.D.K. and P.W. provided assistance with Microarray or RNA sequencing data processing and analysis. L.T., A.M. and G.N. analyzed and interpreted the data, produced the figures, and wrote the manuscript. E.I-B., H.V. and F.V.B. provided conceptual assistance and logistical support. All authors have read and approved the final manuscript.

## Data availability

The data that support the findings of this study are available from the corresponding author upon reasonable request.

## SUPPLEMENTARY FIGURES

**Supplementary Figure S1.**
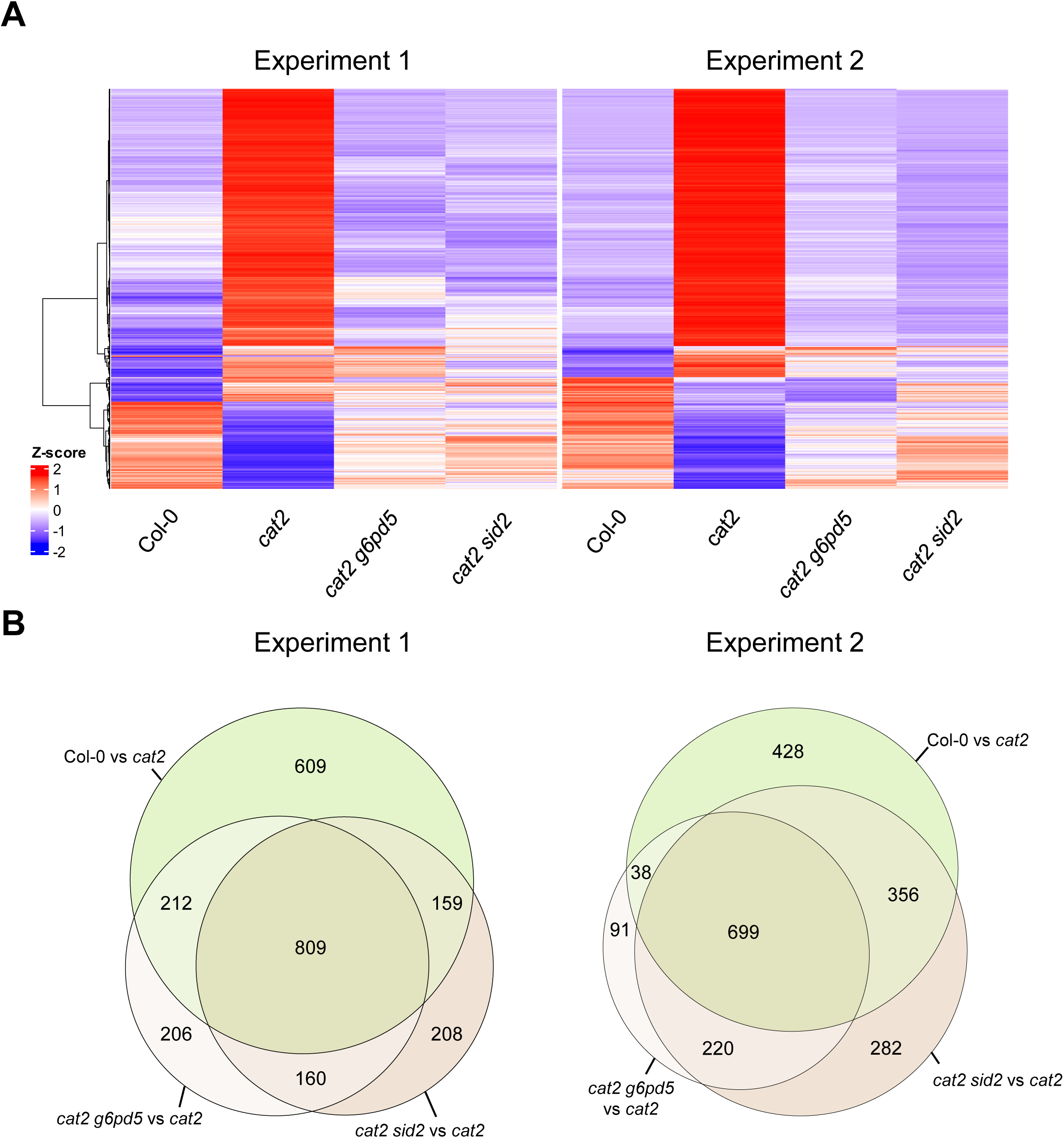
Comparison of expression profiles and differentially expressed genes in *cat2 g6pd5* and *cat2 sid2*. A. Heatmap displaying expression profiles in Col-0, *cat2 sid2*, *cat2 g6pd5* for both experiments. Displayed genes are the 734 *cat2* vs Col-0 DEGs common to both experiments, shown as Z-scores. The top and bottom dendrogram correspond to Pearson correlation with Ward.D2 distance. Z-scores were calculated independently in the two data sets for each gene. B. Venn diagrams showing overlap of differentially expressed genes (DEGs) for the three contrasts comparing *cat2* with Col-0, *cat2 sid2* or *cat2 g6pd5*. The figures indicate the number of DEGs within each specific set. For example, in experiment 1 shown on the left, 609 genes were DEGs between Col-0 and *cat2* that did not also show differential expression between *cat2* and *cat2 sid2* or *cat2* and *cat2 g6pd5*, while 809 genes were found to be differentially expressed in all three comparisons.

**Supplementary Figure S2.**
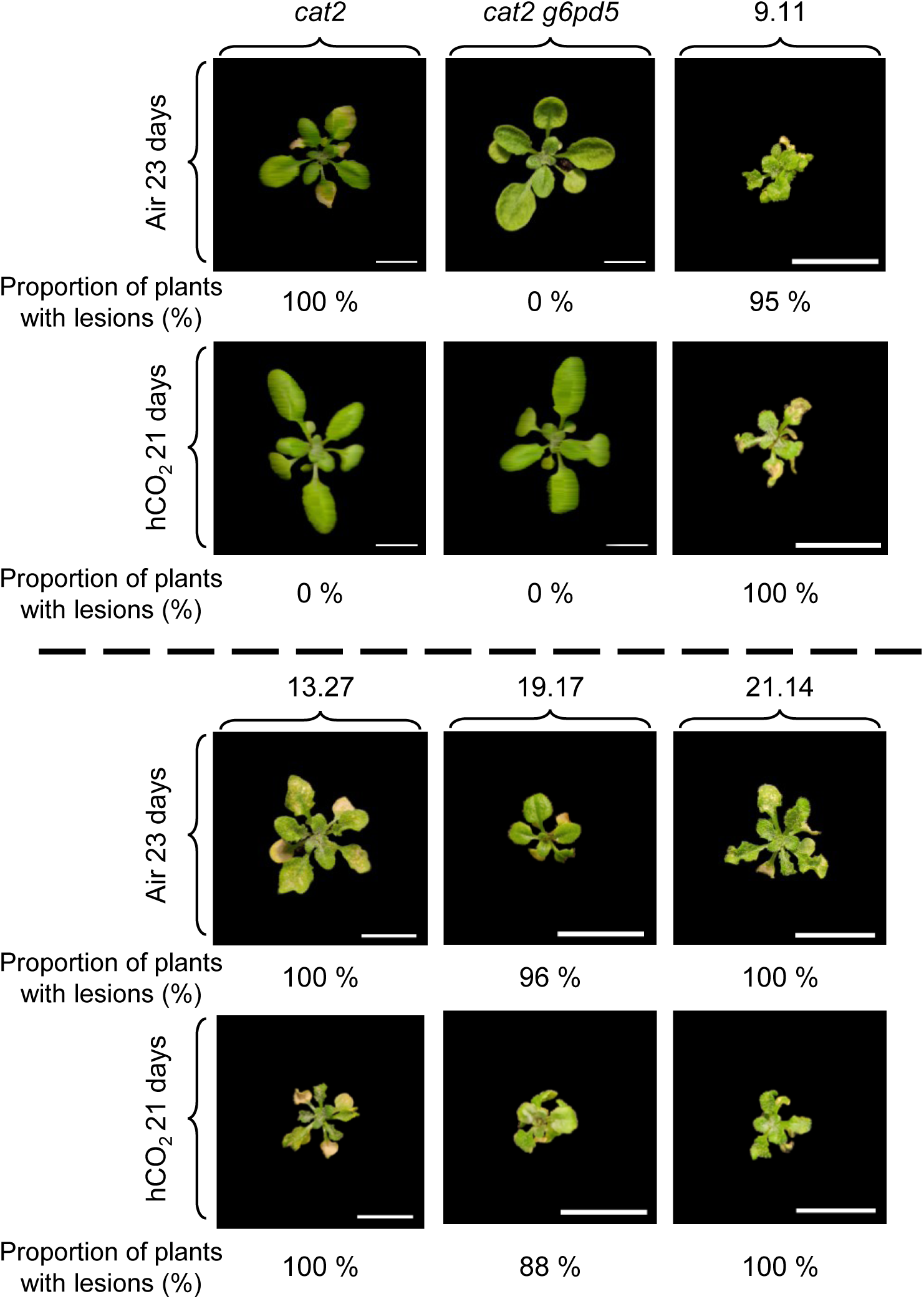
Occurrence of lesion phenotype in air and high CO_2_ for the lines isolated from the genetic screen. Representative phenotype of the plants with lesions after 23 days of growth in air (400 ppm CO_2_) or 21 days of growth at high CO_2_ (3000 ppm CO_2_). If the line does not show a lesion phenotype, a representative plant without lesions is present. Numbers below each picture indicate the percentage of plants presenting lesions (at least 30 plants in each case). Images are presented at different magnifications to clearly display lesion morphology. Scale bars correspond to 1 cm. To allow direct comparison of phenotypes, photographs of reference genotypes (*cat2* and *cat2 g6pd5*) are the same as Fig. 3A.

**Supplementary Figure S3.**
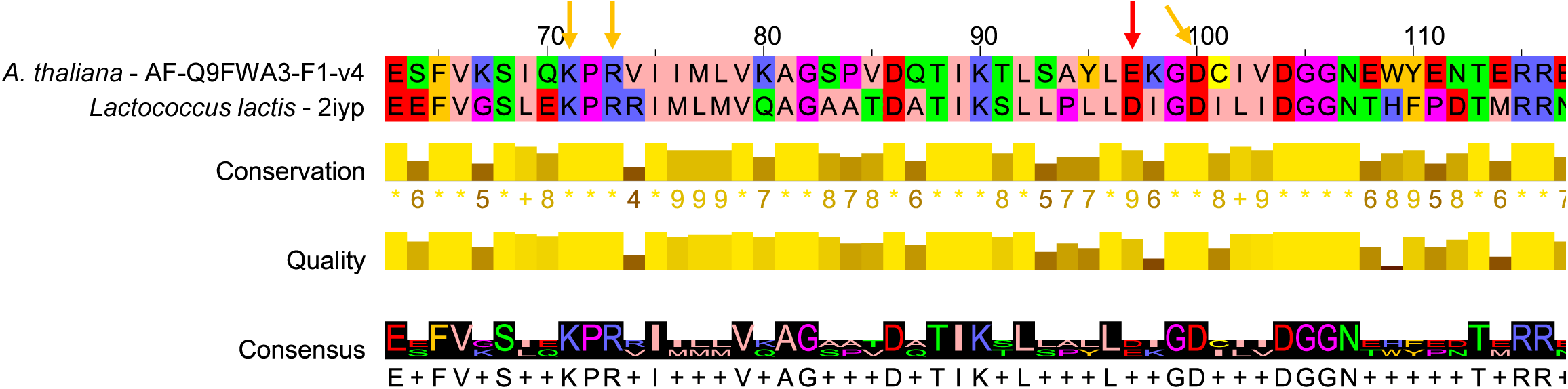
Sequence alignment for *Arabidopsis thaliana* PGD2 and *Lactococcus lactis* GND. Protein sequence alignment of *A. thaliana* PGD2 with *Lactococcus lactis* GND (P96789). The number 1 residue corresponds to the *A. thaliana* proteins. Amino acids are colored according to Zappo colors using Jalview (salmon: Aliphatic/hydrophobic, orange: aromatic, blue: positive, red: negative, green: hydrophilic, pink: conformationally special, yellow: cysteine). Red arrows indicate the residue affected by the SNP (E97 in the wild-type sequence). Orange arrows correspond to residues with hypothetical interactions with E97 according to the AF-Q9FWA3-F1 and Alphafold multimer structures of PGD2.

**Supplementary Figure S4.**
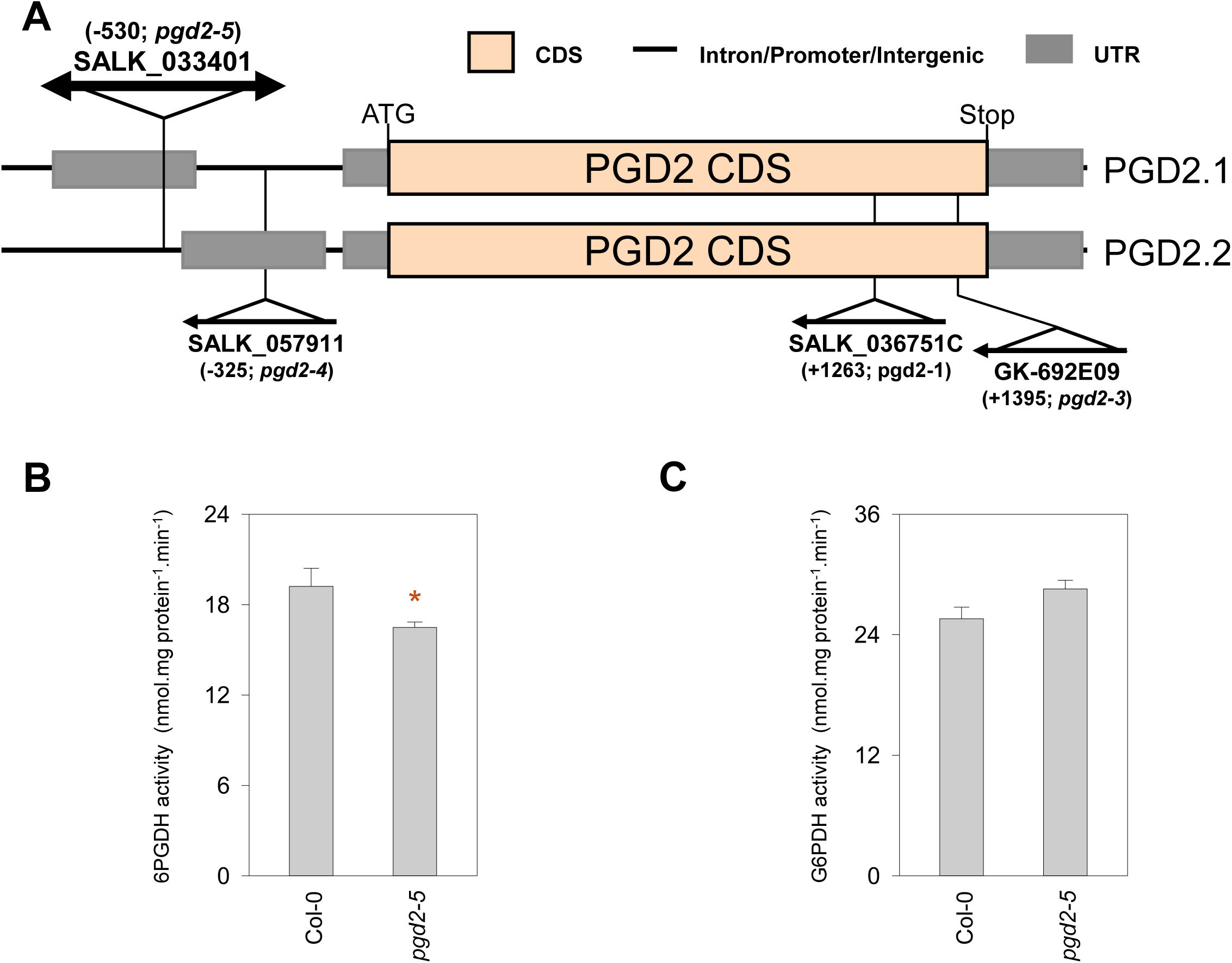
T-DNA insertion and impact of *pgd2-5* allele on 6PGDH and G6PDH enzyme activities. A. Scheme of the T-DNA insertion in the *PGD2* locus for *pgd2-1*, *pgd2-3*, *pgd2-4* & *pgd2-5* alleles depicting the location on the two alternative transcripts of the gene. B. 6PGDH extractable activity in Col-0 and *pgd2-5*. C. G6PDH extractable activity in Col-0 and *pgd2-5*. Analyses were performed on leaf extracts of plants grown from seed for 18 days in 3000 ppm CO_2_ (high CO_2_) and then 8 days in air (400 ppm CO_2_). Data are means ± SE of three biological replicates. Significant differences relative to Col-0 are indicated by + (Student’s t-test, p < 0.05).

**Supplementary Figure S5.**
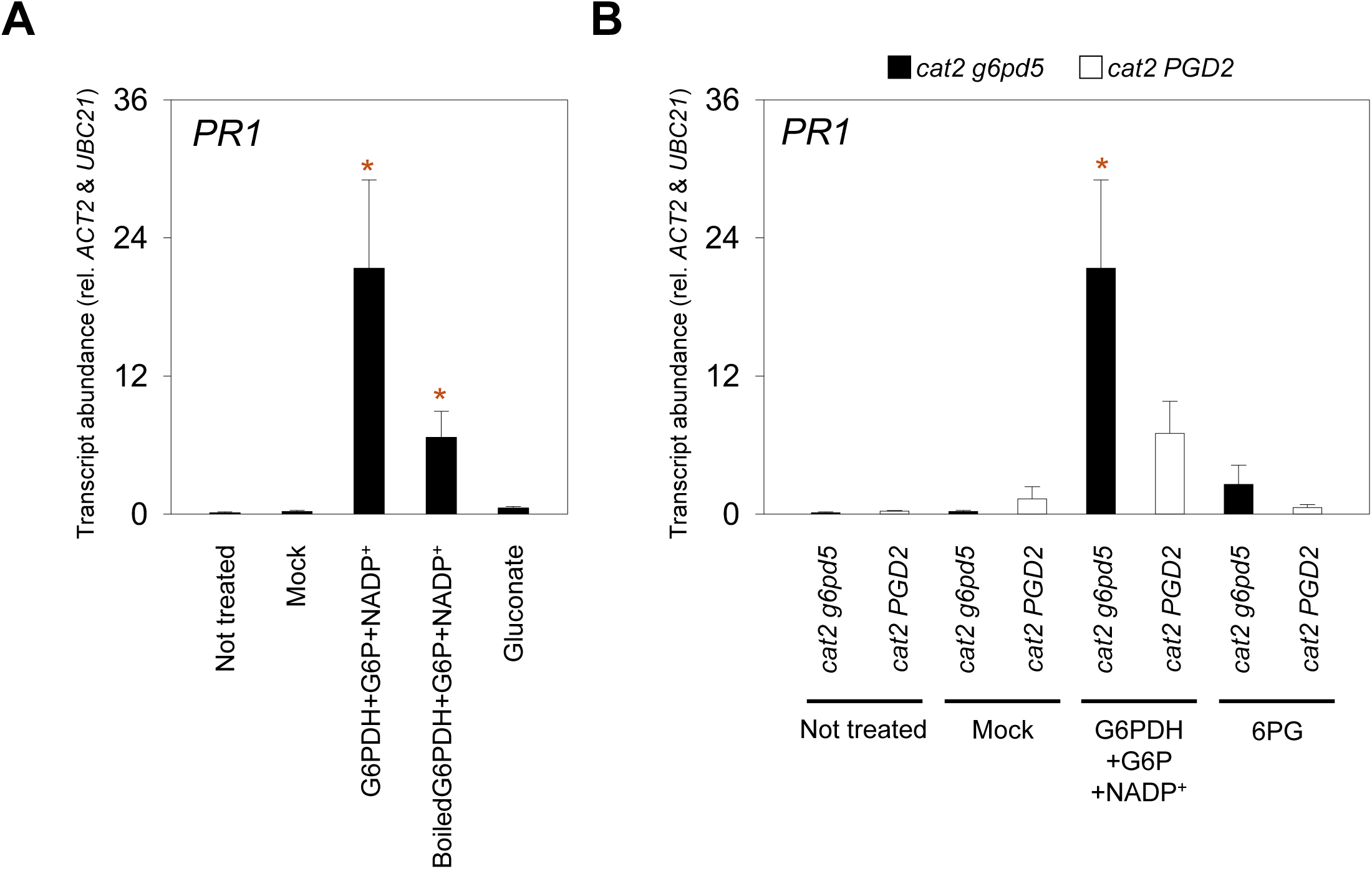
Impact of treatment investigating a role for a signal metabolite related to the OPPP. A. SA marker gene *PR1* abundance 24 h after infiltration of the second leaf pairs of *cat2 g6pd5*. B. SA marker gene *PR1* abundance 24 h after infiltration of the second leaf pairs of *cat2 g6pd5* (black) and *cat2* PGD2 (grey). Sampled plants were grown for 22 days in air and long days at 200 µmol photon/m²/s. G6PDH: G6PDH from yeast, 5 U/mL; G6P: glucose 6-phosphate, 15 mM; NADP_+_: 600 µM; 6PG: 6-phosphogluconate, 15 mM; Gluconate, 15 mM. Boiled G6PDH indicates G6PDH solution incubated for 15 min at 95 °C and then transferred to 4°C before mixing with other compounds. Activity of an aliquot of the boiled mixture was measured by following NADPH production at A340 to confirm inactivation compared to G6PDH. The *cat2* PGD2 genotype corresponds to the line *cat2* PGD2 T3-1. Transcript abundance is normalized on *UBC21* and *ACT2* housekeeping gene levels. Data are means ± SE of three biological replicates. Significant differences relative to mock treatments are indicated by * (Student’s t-test, p < 0.05).

---

**Supplementary Table S1:** List of primers used in this study

**Supplementary Table S2:** Microarray data.

**Supplementary Table S3:** RNA-seq data.

